# *EBSn,* a robust synthetic reporter for monitoring ethylene responses in plants

**DOI:** 10.1101/2025.05.23.655144

**Authors:** Josefina-Patricia Fernandez-Moreno, Mario Fenech, Anna E. Yaschenko, Chengsong Zhao, Matthew Neubauer, Hannah N. Davis, Alex J. Marchi, Raine Concannon, Alexandra Keren-Keiserman, Moshe Reuveni, Victor Levitsky, Dmitry Oshchepkov, Vladislav Dolgikh, Alexander Goldshmidt, José T. Ascencio-Ibáñez, Elena Zemlyanskaya, Jose M. Alonso, Anna N. Stepanova

## Abstract

Ethylene is a gaseous plant hormone that controls a wide array of physiologically relevant processes, including plant responses to biotic and abiotic stress, and induces ripening in climacteric fruits. To monitor ethylene in plants, analytical methods, phenotypic assays, gene expression analysis, and transcriptional or translational reporters are typically employed. In the model plant Arabidopsis, two ethylene-sensitive synthetic transcriptional reporters have been described, *5xEBS:GUS* and *10x2EBS-S10:GUS*. These reporters harbor a different type, arrangement, and number of homotypic *cis*-elements in their promoters and thus may recruit the ethylene master regulator EIN3 in the context of alternative transcriptional complexes. Accordingly, the patterns of GUS activity in these transgenic lines differ and neither of them encompasses all plant tissues even in the presence of saturating levels of exogenous ethylene. Herein, we set out to develop and test a more sensitive version of the ethylene-inducible promoter that we refer to as *EBSnew* (abbreviated as *EBSn*). *EBSn* leverages a tandem of ten non-identical, natural copies of a novel, dual, everted, 11bp-long EIN3-binding site, *2EBS(−1)*. We show that in Arabidopsis, *EBSn* outperforms its predecessors in terms of its ethylene sensitivity, having the capacity to monitor endogenous levels of ethylene and displaying more ubiquitous expression in response to the exogenous hormone. We demonstrate that the *EBSn* promoter is also functional in tomato, opening new avenues to manipulating ethylene-regulated processes, such as ripening and senescence, in crops.

## Introduction

Plant hormone ethylene is a multi-functional growth regulator involved in plant responses to biotic and abiotic stresses and in the control of a wide array of developmental processes, from seed germination to organ senescence and fruit ripening (Matoo, 1991; Abeles et al. 1992).

Mechanistic understanding of how and where endogenous ethylene is produced and acts in a plant opens the possibility of modulating a myriad of ethylene-regulated processes for greater plant resilience and extended produce shelf life (Li et al. 2019; Wang et al. 2020). Studies in Arabidopsis and other plant species have uncovered the key molecular components of ethylene biosynthesis and signaling pathways (Johnson and Ecker, 1998; Fernandez-Moreno and Stepanova, 2019; Binder 2020). Ethylene is produced from the amino acid methionine via 1-aminocyclopropane-1-carboxylic acid (ACC), with the rate-limiting enzymes ACC synthase and ACC oxidase catalyzing the two final steps leading to ethylene production. Ethylene is a gaseous signal perceived by endoplasmic reticulum (ER) membrane-localized receptors that in Arabidopsis are represented by five family members: ETHYLENE RESPONSE1 (ETR1), ETR2, ETHYLENE INSENSITIVE4 (EIN4), ETHYLENE RESPONSE SENSOR (ERS1) and ERS2 (Chang et al. 1993; Hua et al., 1995 and 1998; Sakai et al., 1998; Gao et al. 2008). Binding of ethylene turns the receptors off, resulting in the inactivation of a Raf-like kinase CONSTITUTIVE TRIPLE RESPONSE1 (CTR1) (Kieber et al. 1993; Park et al. 2023) on the cytoplasmic surface of the ER and the dephosphorylation of a positive regulator of ethylene signaling, EIN2, an ER-localized, transmembrane protein (Alonso et al. 1999). The C-terminus of the dephosphorylated EIN2, cleaved in the presence of ethylene by an unknown protease, plays cytoplasmic and nuclear roles in ethylene signaling. In the cytoplasm, it inhibits the translation of transcripts for EIN3-BINDING F-BOX1 (EBF1) and EBF2 proteins (Li et al. 2015; Merchante et al. 2015). EBFs, in turn, are responsible for the proteasomal turnover of transcriptional master regulators of ethylene signaling, EIN3 and EIN3-LIKE (EILs), in the absence of ethylene (Song et al., 2015; Fernandez-Moreno et al., 2019; Binder 2020). In the presence of ethylene, *EBF*s are not efficiently translated, EIN3/EILs are stabilized and initiate a transcriptional cascade of events by binding to the promoters of ethylene-regulated target genes. The transcriptional activity of EIN3 requires the function of the EIN2 C-terminus in the nucleus (Alonso et al. 1999; Ju et al. 2012; Wen et al. 2012). The mechanism of this regulation is not fully understood but likely involves chromatin structure modification (Wang et al. 2021).

EIN3 and EIL proteins are thought to primarily work by activating target gene transcription (Dolgikh et al., 2019; Binder 2020), but they have also been implicated in the repression of a subset of target genes, a function accomplished by complexing with TRANSCRIPTIONAL REPRESSOR OF EIN3-DEPENDENT ETHYLENE-RESPONSE1 (TREE) proteins (Wang et al. 2020). EIN3/EILs are thought to bind to gene promoters in different configurations dictated by the architecture and accessibility of target promoter elements (Dolgikh et al., 2019). The consensus EIN3/EIL binding site (EBS) inferred from *in vitro* and *in vivo* studies in Arabidopsis is *ATGTAT* (Solano et al. 1998; Chang et al. 2013; Song et al. 2015; O’Malley et al. 2016). In the promoters of genes bound by EIN3 in DAP-seq (O’Malley et al. 2016) and ChIP-seq (Chang et al. 2013) datasets, this and related elements can be present alone or in tandems (in direct, inverted, or everted orientations). Accordingly, some gene targets of EIN3/EIL are thought to be compatible with EIN3/EIL binding to DNA as monomers (or, potentially, as dimers with other EIN3/EILs or sequence-unrelated proteins), whereas other promoters are likely to recruit EIN3/EIL in a dimeric form (with or without other interactors). Thus, it should be possible to leverage these different configurations of EIN3/EIL binding sites (EBSs) to capture the activity of alternative EIN3/EILs complexes (Fernandez-Moreno et al. 2024). By building different versions of synthetic EBS promoters, it should be possible to create divergent transcriptional reporters for monitoring ethylene responses mediated by EIN3/EIL1 present in specific transcriptional complexes.

We have previously generated reporters with two types of synthetic ethylene promoters, *5xEBS* and *10x2EBS-S10* (Stepanova 2001; Fernandez-Moreno et al. 2024). The promoter in the *5xEBS:GUS* construct consists of a *35S(−46)* minimal promoter, preceded by five direct copies of a natural EIN3-binding element derived from the promoter of an Arabidopsis ethylene-inducible gene, *ETHYLENE RESPONSE DNA-BINDING FACTOR1* (*EDF1*) (Stepanova 2001; Alonso et al. 2003; Stepanova at al. 2007) (**Fig. 1, Supplementary Fig. S1A**). The promoter in *10x2EBS-S10:GUS* consists of the same *35S(−46)* core preceded by 10 copies of a synthetic dual everted *EBS* element separated by an A-rich 10nt spacer (**Fig. 1, Supplementary Fig. S1B**) (Fernandez-Moreno et al. 2024). The *EDF1* promoter is a direct target of EIN3, as demonstrated by gel-shift (Stepanova 2001) and ChIP-seq (Chang et al. 2013) assays, whereas the *2EBS-S10* sequence is a synthetic DNA element optimized for EIN3 binding using gel-shift assays (Song et al. 2015).

**Figure 1.**
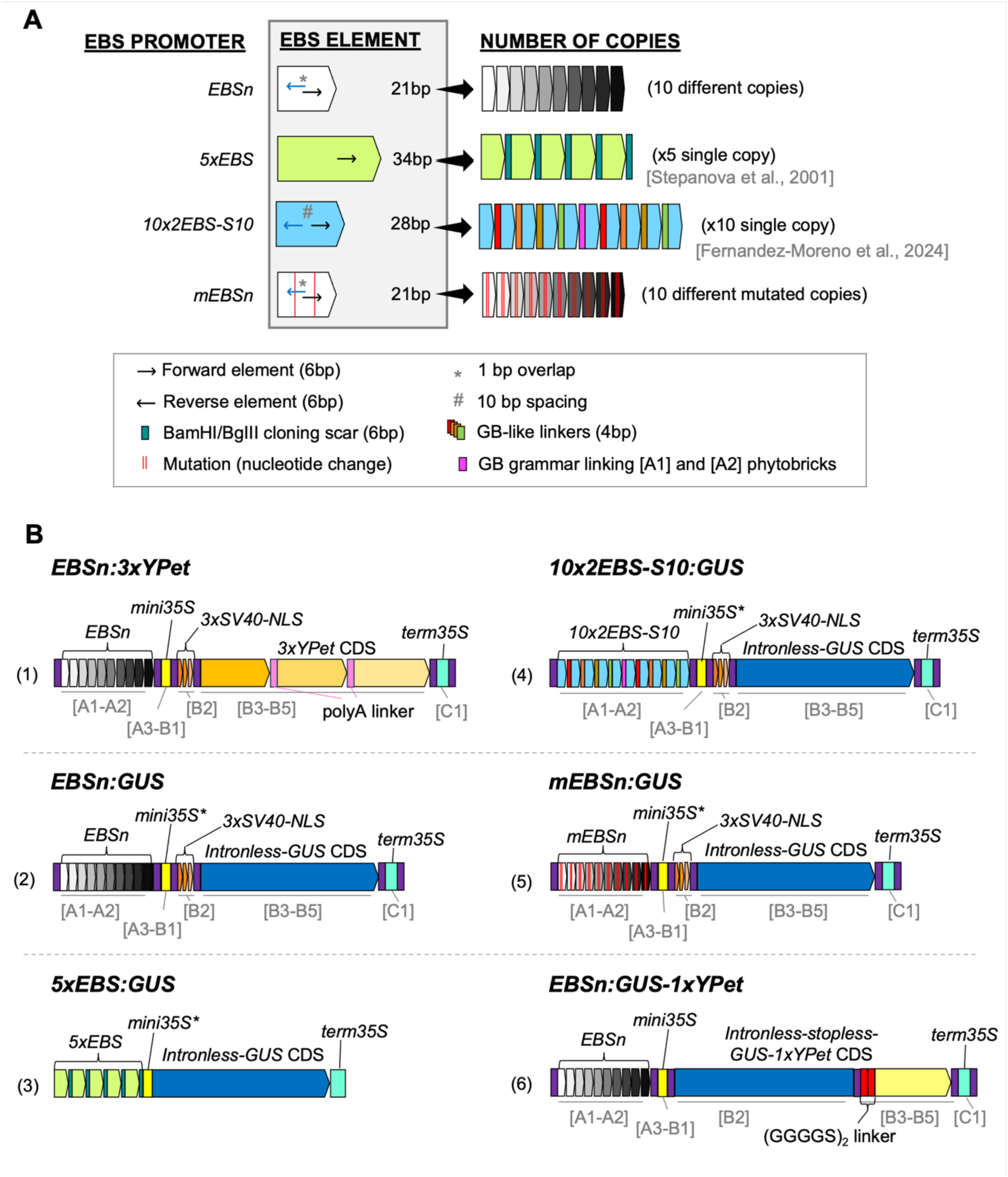
Graphical representation of ethylene-inducible proximodistal promoters and transcriptional reporters utilized in this work. **A)** Simplified representation for the size and number of *EIN3 binding site* (*EBS*) elements within different ethylene-inducible promoters experimentally tested in this study. Two of the *EBS* versions have previously been described, *5xEBS* (Stepanova et al 2001) and *10x2EBS-S10* (Fernandez-Moreno et al. 2024), and two were generated and characterized in this work, *EBSnew* (*EBSn*) and mutated*EBSnew* (*mEBSn*). **B)** Schematics of the transcriptional reporters harboring the *EBS*-containing promoters shown in panel A. For every reporter but (3), the GoldenBraid (GB) grammar for each assembled phytobrick is represented in purple at the flanks and between DNA parts marked with square brackets beneath the graphical representations. Reporters (1), (2), (4) and (5) harbors five phytobricks: [A1-A2] distal promoter (*EBSn*, *10x2EBS-S10* or *mEBSn*), [A3-B1] core promoter (*mini35S*/*mini35S\**), [B2] nuclear localization signal (*3xSV40-NLS*), [B3-B5] reporter CDS (*3xYPet* or *intronless--GUS*), and [C1] terminator (*term35S*) (**Supplementary Table S1A**). Reporter (6) lacks a subcellular localization signal (no *3xSV40-NLS*) and harbors a [B2-B5] *GUS-1xYPet* double reporter assembled from a [B2] *intronless-stopless--GUS* CDS and a [B3-B5] *(Gly-Gly-Gly-Gly-Ser)_2_ linker-1xYPet* phythobricks (**Supplementary Table S1A**). Finally, reporter (3) represents the original *5xEBS* reporter (Stepanova 2001) which harbors a distal promoter with five *EBS* elements in tandem spaced by a BamH/BgIII cloning scare, a core promoter (*mini35S\**), an *intronless GUS* CDS reporter, and a terminator (*term35S*). Core promoters *mini35S* (54bp long) and *mini35S\** (60bp long) have 49bp in common, with variable 5bp flanks (**Supplementary Fig. S1C**).

The two respective *GUS* reporters, *5xEBS:GUS* and *10x2EBS-S10:GUS,* showed clearly divergent patterns of expression in seedlings and inflorescences (Fernandez-Moreno et al. 2024), supporting the idea that the architecture of EIN3 target sites in the promoters dictates the patterns of reporter expression, presumably by selectively recruiting EIN3/EIL proteins existing in certain protein complexes (with itself and other transcription factors (TFs)) while discriminating against EIN3/EILs present in other configurations incompatible with the specific binding site architecture. Importantly, neither the *5xEBS* nor the *10x2EBS-S10* promoter is active in all seedling tissues, even though in ethylene-treated etiolated seedlings, most, if not all, cells appear to respond to exogenous ethylene phenotypically (Fernandez-Moreno et al. 2024), suggesting that the two synthetic promoters are not capturing all possible EIN3/EIL configurations present in plants.

Herein, we describe a novel ethylene-inducible promoter, *EBSnew* (abbreviated as *EBSn*), with a distinct *EBS* element architecture (**Fig. 1A, Supplementary Fig. S2A**). We show that in Arabidopsis this promoter is more sensitive to endogenous ethylene than *5xEBS* or *10x2EBS-S10* and that it becomes broadly activated across different seedling tissues in response to exogenous ethylene. We demonstrate that the high basal activity of the *EBSn* promoter in seedlings is *EIN2*-dependent and thus ethylene-mediated, and that the promoter is also active in adult Arabidopsis plants, as well as in tomato. We propose future applications of this promoter for controlling ethylene-regulated processes to delay produce deterioration pre- and post-harvest and to reduce food waste.

## Results and Discussion

### The *EBSn* promoter responds to ethylene in Arabidopsis seedlings

The *EBSn* promoter consists of a tandem of 10 non-identical copies of a dual everted EIN3 binding site (where the two half-sites are positioned with a one base pair (bp) overlap with respect to one another, and each 11bp-long dual *EBS* site is separated by a 10 nucleotide spacer) placed upstream of a minimal *35S(−46)* promoter (**Fig. 1A, Supplementary Fig. S2A**). To test the functionality of this composite synthetic *EBSn* distal promoter, we first generated fusions with a *35S(−46)* core promoter, a triple yellow fluorescent protein gene *3xYPet*, a viral nuclear localization signal *SV40,* and a *35S* terminator (**Fig. 1B, Supplementary Table S1A**), and transformed the reporter into wild-type Arabidopsis plants (Columbia-0). Over 50 T1 lines were generated and two single-insertion lines, referred to as A and C, were examined in detail. In three-day-old dark-grown seedlings, Arabidopsis lines harboring this transgene showed low but clearly detectable levels of basal expression in the apical hooks and root tips in control air conditions (**Fig. 2**). The expression was strongly and ubiquitously induced in response to 10 ppm exogenously applied ethylene (**Fig. 2A, B**) or saturating concentrations of the ethylene precursor ACC (10 μM) (**Supplementary Fig. S3**). In both cases the strongest fluorescence was detected in the apical hooks, cotyledons, root tips, root-hypocotyl junctions, and vasculature (**Fig. 2, Supplementary Fig. S3**). This expression pattern was maintained faithfully without any silencing across multiple generations (at least until T5). As shown for propidium iodide-stained roots of ACC-treated seedlings, the reporter is predominantly detected in the nuclei of the root cap, epidermis, and columella cells (**Fig. 2C, D**). Remarkably, as little as 10 nM ACC was sufficient to trigger a prominent expansion of the *EBSn:3xYPet* expression domain in the root tip (**Supplementary Fig. S4**), indicating the high sensitivity of the reporter.

**Figure 2.**
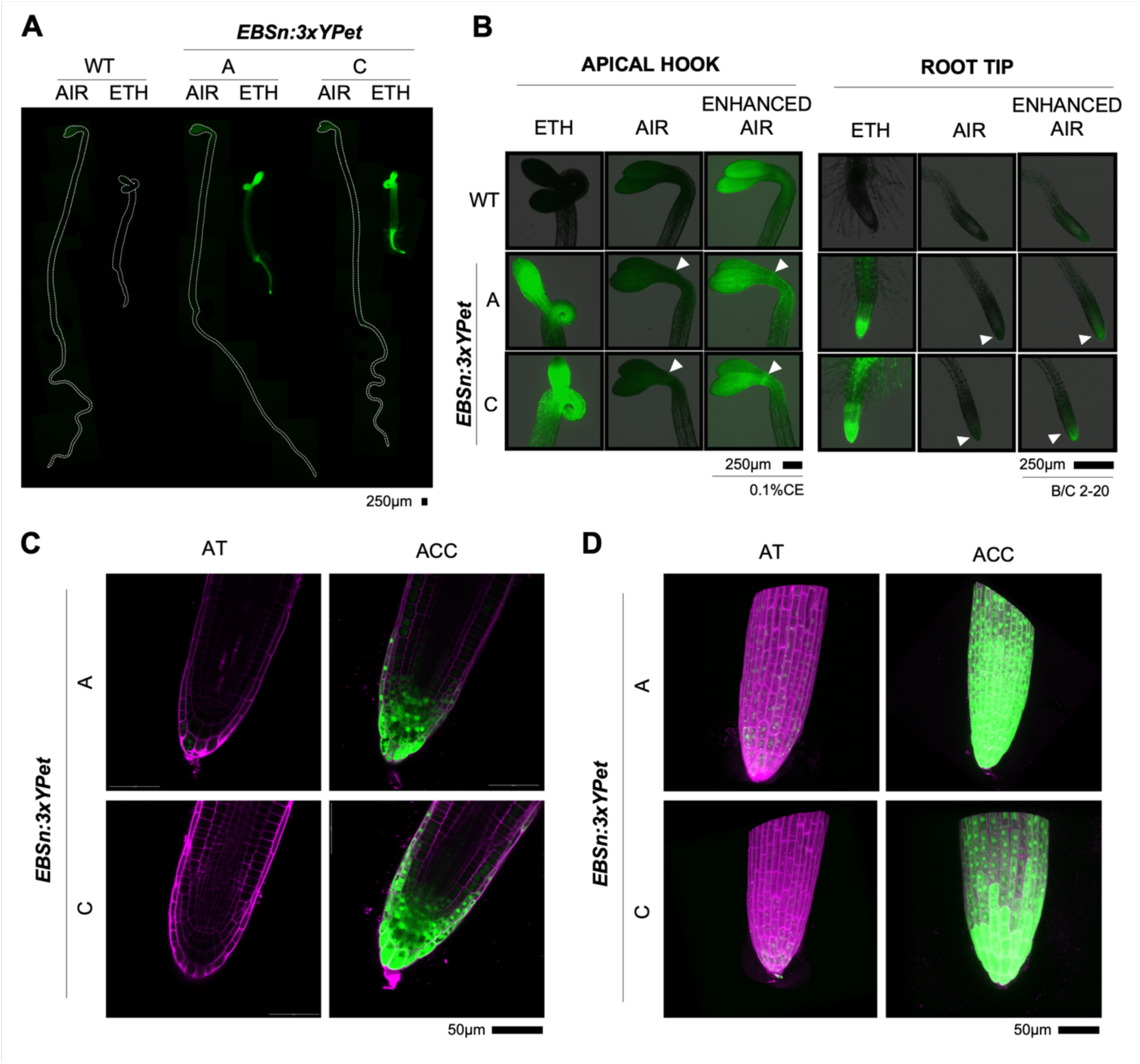
The *EBSn:3xYPet* reporter is induced by ethylene and ACC in three-day-old dark-grown Arabidopsis seedlings. **A)** Whole-plant fluorescence images of wild-type (WT) and transgenic (*EBSn:3xYPet,* lines A and C) seedlings germinated in AT media in the presence of hydrocarbon-free air (AIR) or 10 ppm ethylene (ETH). **B)** Close-up apical hook and root tip images of seedlings from panel A. AIR and ETH images are shown, along with ENHANCED AIR images that were digitally enhanced (0.1% contrast enhancement, CE, for apical hooks and 2-20 min-max brightness/contrast, B/C, for root tips in ImageJ v1.52, Fiji 2015) to show nuclear localization of 3xYPet in apical hooks and a more diffuse 3xYPet signal in root tips (arrowheads) not seen in WT plants. **C)**, **D)** Confocal microscopy images displaying optical cross-sections (C) and Z-stacks (D) of propidium iodide-stained (pink) root tips of *EBSn:3xYPet* seedlings (lines A and C) germinated in AT media or AT media supplemented with 10 μM ACC.

Given the recently uncovered ethylene-independent roles of ACC in plants (Vanderstraeten et al. 2019; Li et al. 2020; Mou et al. 2020 and 2025), we tested whether the inducibility of the *EBSn:3xYPet* reporter by ACC requires ethylene signaling by crossing this reporter to a strong ethylene insensitive mutant *ein4* that carries a missense gain-of-function mutation (Hua et al. 1998). The dominant nature of this mutant enabled us to run the experiment in the F1 generation of the cross. The *ein4* mutation was able to greatly suppress the response of *EBSn:GUS* to ACC (**Supplementary Fig. S5A**), confirming that intact ethylene signaling is required for this reporter’s responsiveness to ACC and implying that the reporter responds to ACC-derived ethylene. The residual activity of the reporter in the cotyledons and root tips of *ein4/+* heterozygous plants in ACC suggests that this mutant may not be fully dominant or that *EIN4* may not be ubiquitously expressed at adequate levels for its dominant mutant version to fully block ethylene signaling in all tissues. Alternatively, *EBSn* may also respond to signals other than ethylene or the minimal *35S(−46)* promoter may have basal activity on its own.

To further test the ethylene dependency of the *EBSn:3xYPet* activity, we next crossed *EBSn:3xYPet* with a recessive strong ethylene insensitive mutant, *ein2-5*, that is caused by a 7-nucleotide coding region deletion (Alonso et al. 1999) and analyzed the F4 plants homozygous for both *ein2-5* and the reporter (**Fig. 3**). Much like with *ein4/*+ plants, the *ein2* mutation was able to block a majority of the *EBSn:3xYPet* signal, consistent with the idea that nearly all of the *EBSn* promoter activity is ethylene-mediated. However, in one of the two transgenic lines, line A (but not in line C), very low levels of residual fluorescence were still detected in the root tip when the fluorescence signal was digitally enhanced (**Supplementary Fig. S6**). However, since this signal was not dependent on ethylene treatment, we conclude that this background activity is not ethylene-regulated and may be a consequence of basal reporter expression from the minimal *35S(−46)* promoter and/or enhancer-like effects of the genomic regions flanking the transgene.

**Figure 3.**
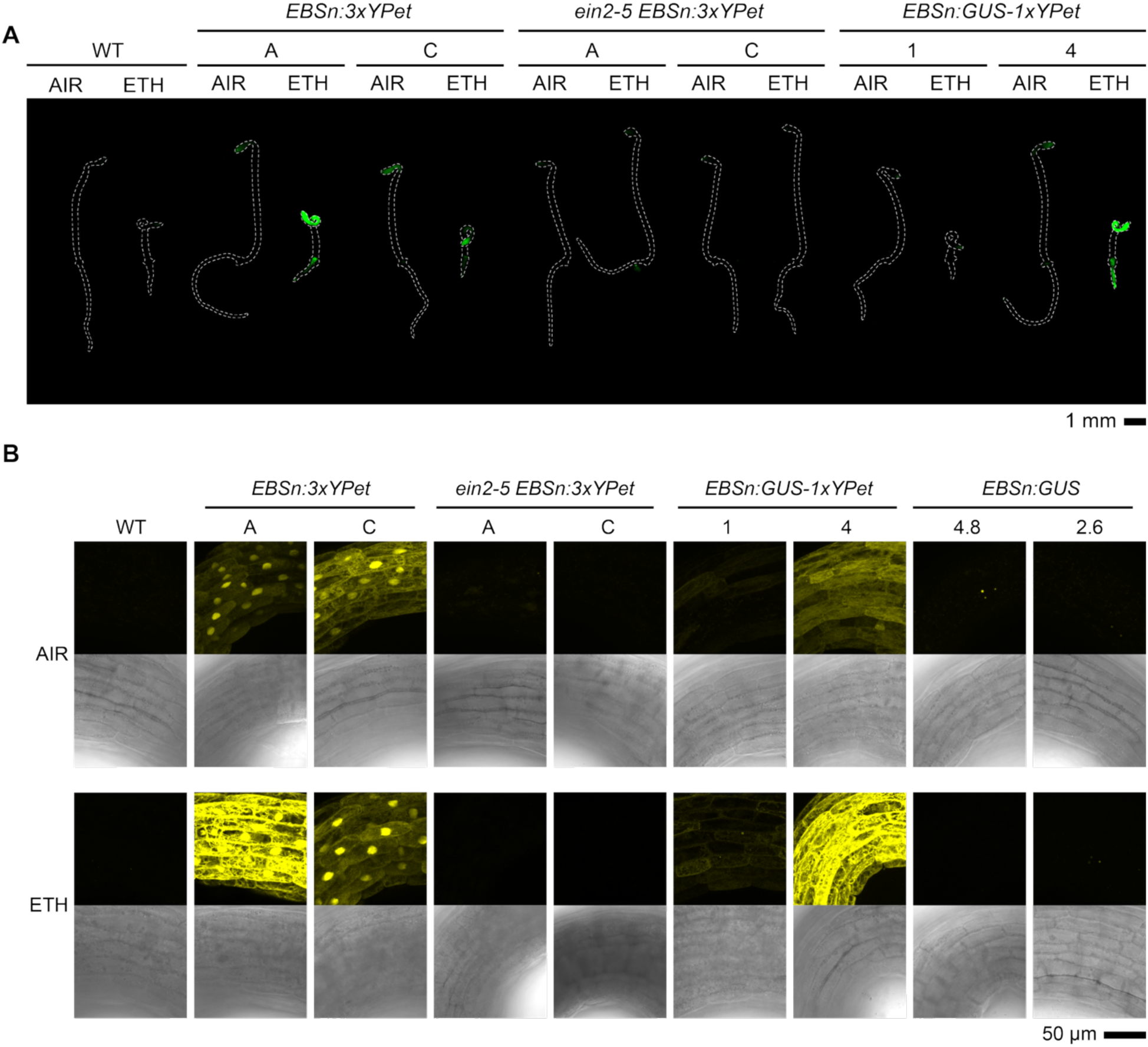
Robust fluorescence of *EBSn*-driven reporters in three-day-old dark-grown Arabidopsis seedlings is ethylene-mediated. Seeds were germinated in AT plates in the presence of hydrocarbon-free air (AIR) or 10 ppm ethylene (ETH). **A)** The loss of functional *EIN2* in *ein2-5 EBSn:3xYPet* (lines A and C) abolishes the ethylene-dependent triple response and YPet activity in 10 ppm ethylene, and the patterns of YPet fluorescence in *EBSn:GUS-1xYPet* dual reporter lines 1 (weak) and 4 (strong) are comparable to that of the *EBSn:3xYPet* lines A and C. **B)** Confocal images of the *EBSn* activity in the apical hooks of three-day-old dark-grown *EBSn:3xYPet* and *EBSn:GUS-1xYPet* seedlings in Arabidopsis. Seeds were germinated in AT plates in the presence of hydrocarbon-free air (AIR) or 10 ppm ethylene (ETH). Fluorescence and bright-field images are displayed.

Since this basal activity was very weak (i.e., not detectable without digital image enhancement) and only observed in one of the two lines of *ein2-5 EBSn:3xYPet* (**Supplementary Fig. S6**), we conclude that both transgenic lines of the *EBSn:3xYPet* reporter are suitable for monitoring seedling responses to endogenous or exogenous ethylene.

We also generated a *GUS* version of the *EBSn* reporter. The *EBSn* proximodistal promoter was again fused with the *35S(−46)* core promoter driving *GUS* followed by a *35S* terminator (**Fig. 1B, Supplementary Table S1A**). This construct enabled us to compare the performance of *EBSn:GUS* with that of the classic *5xEBS:GUS* and *10x2EBS-S10:GUS* reporters we developed previously (Stepanova 2001; Stepanova et al. 2007; Fernandez-Moreno et al. 2024). As with the *3xYPet* version of the *EBSn* reporter, we generated over 30 T1 lines, but focused our efforts on characterizing two of the lines, a moderately expressing line (4.8) and a stronger line (2.6).

Under control (air) conditions, the majority of the *EBSn:GUS* lines, exemplified by line 4.8, showed strong expression in the apical hooks, upper hypocotyls, cotyledons, and root tips (**Fig. 4A**), suggesting that the *GUS* reporter can detect endogenous levels of ethylene with greater sensitivity relative to its *3xYPet* counterpart. In a few stronger lines, exemplified by line 2.6, GUS activity expanded to lower hypocotyls and the root (**Fig. 4A**). Noteworthy, in the presence of an ethylene receptor inhibitor 1-methylcyclopropene (1-MCP, 200 ppm), the activity of *EBSn:GUS*, although clearly reduced, was not fully eliminated, as shown for a moderate line 4.8, or was largely ineffective, as demonstrated for the stronger line, 2.6 (**Fig. 4A**). These results suggest that 1-MCP may not be able to fully block the ethylene response despite clearly enhancing etiolated seedling elongation that is indicative of ethylene response inhibition (**Fig. 4A**).

**Figure 4.**
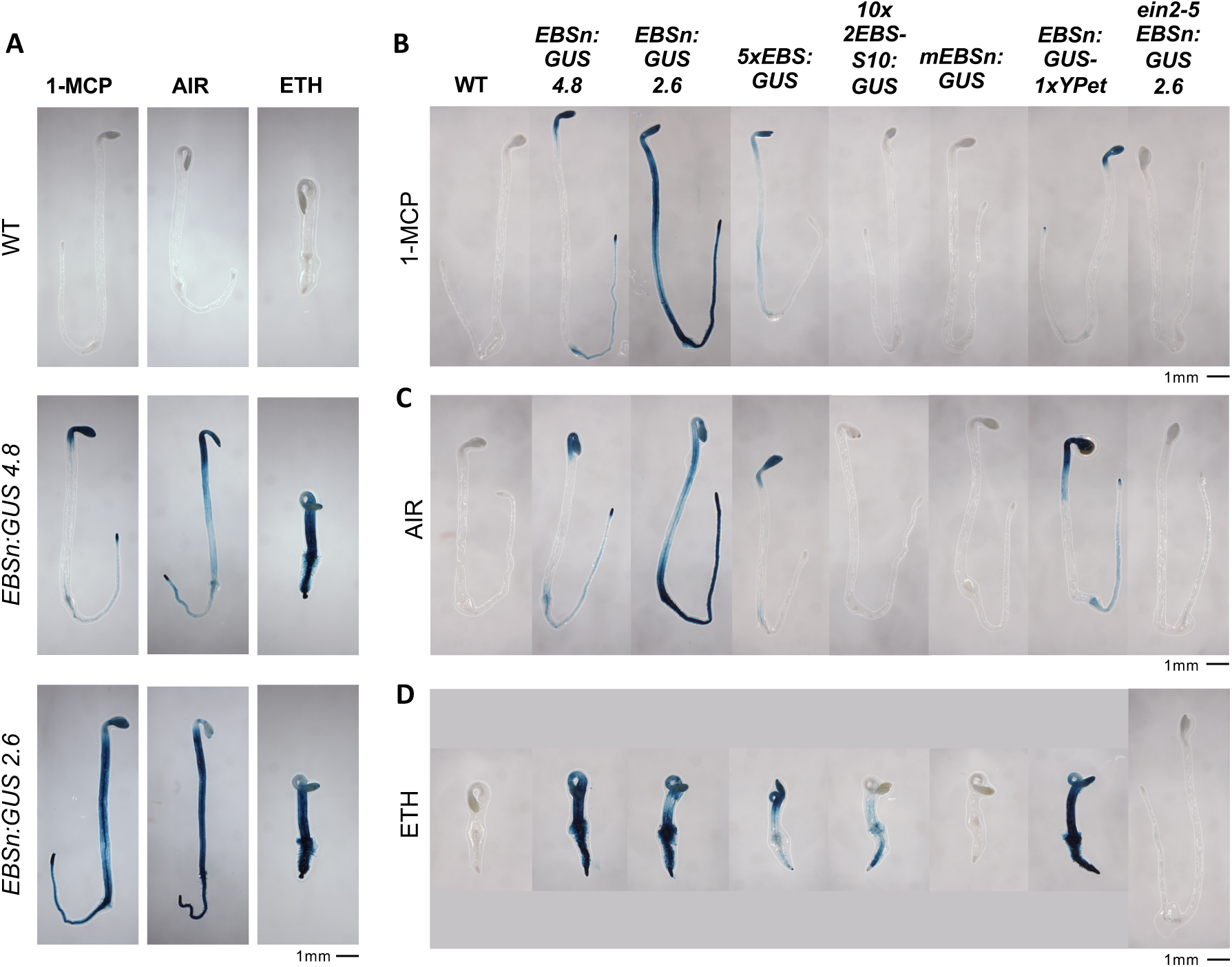
*EBSn* is more uniformly activated by ethylene in Arabidopsis seedlings than are other synthetic ethylene-regulated *EBS* promoters. Three-day-old seedlings germinated in AT plates in the dark for three days in the presence of 200 ppm 1-MCP (ethylene receptor inhibitor), hydrocarbon-free air (AIR), or 10 ppm ethylene (ETH) were stained for GUS overnight and photographed. **A)** Expression of *EBSn:GUS* in representative moderate (4.8) and strong (2.6) lines. **B), C), D)** A side-by-side comparison of *GUS* expression in *EBSn:GUS* (lines 4.8 and 2.6)*, 5xEBS:GUS, 10x2EBS-S10:GUS, mEBSn:GUS, EBSn:GUS-1xYPet, ein2 EBSn:GUS* (2.6), and WT Col-0 in 1-MCP (panel A), air (panel B) or ethylene (panel C). Scale bars represent 1mm.

As expected, in seedlings germinated in the presence of 10 ppm ethylene (**Fig. 4A**) or 10 μM ACC (**Supplementary Fig. S7**), the activity of the *EBSn:GUS* reporter was dramatically induced, with the GUS staining now encompassing a majority of the seedling in both moderate (4.8) and strong (2.6) *GUS* lines. This staining pattern is consistent with the aforementioned ubiquitous activation of the *YPet* version of the reporter driven by the same promoter (**Fig. 2A, Supplementary Fig. S3**). Thus, both of these reporters are suitable for monitoring the ethylene response in different organs of etiolated seedlings.

In contrast, the two previously published ethylene reporters, *5xEBS:GUS* and *10x2EBS-S10:GUS,* had little to no expression in control air conditions or 1-MCP (**Fig. 4B, C**), suggesting that these older reporters are not sensitive enough to detect low endogenous concentrations of ethylene in most tissues. In the presence of exogenous ethylene (10 ppm), *5xEBS:GUS* was activated predominantly in the apical hooks, cotyledons and root tips, whereas *10x2EBS-S10:GUS* was turned on in the hypocotyls and roots (**Fig. 4D**). The difference in the sensitivity and expression patterns between the three *GUS* reporters is likely to be the consequence of a different *EBS* copy number (which may dictate the amplitude of the transcriptional response) and alternative *EBS* site architecture (which may recruit EIN3 in the context of different TF complexes with different tissue distribution and DNA binding preferences) (Fernandez-Moreno et al. 2024).

### The basal expression of *EBSn:GUS* in Arabidopsis seedlings is ethylene-mediated

Given the prominent expression of *EBSn:GUS* in 1-MCP, we wondered whether the basal levels of GUS activity (despite the high doses of this supposedly very potent ethylene receptor antagonist) are caused by (1) the activity of the minimal *35S(−46)* core promoter, (2) the binding of a TF other than EIN3 to the distal *EBSn* promoter (e.g., to DNA element(s) created at the junction(s) between adjacent copies of the *EBS*), or (3) residual ethylene responses in the presence of 1-MCP. To test these hypotheses, we generated a mutant version of the *EBSn* promoter where a single nucleotide substitution is introduced in every copy of the *EBS* half-site (*mEBSn* in **Fig. 1A, B, Supplementary Fig. S2B, Supplementary Table S1A**). These mutations are expected to abolish EIN3 binding (Solano et al. 1998). No expression was detected in any of the transgenic lines harboring *mEBSn:GUS* either with or without ethylene (**Fig. 4B, C, and D**), indicating that the minimal promoter alone cannot drive the reporter expression in etiolated seedlings and that no additional TF binding (beyond that of EIN3/EILs) to *EBSn* is occurring to support this reporter’s expression. This leaves residual ethylene response as the most likely explanation for the basal activity of the wild-type version of *EBSn:GUS* in 1-MCP.

To further validate the residual ethylene response as the explanation for basal expression of wild-type *EBSn:GUS* in 1-MCP, we crossed the reporter with the aforementioned dominant ethylene insensitive mutant *ein4* and examined the F1 generation of the cross. As in the cross with the *EBSn:3xYPet* construct, *ein4/*+ was able to dramatically reduce but not fully block the *EBSn:GUS* activity in F1s of crosses, with the apical hooks and roots tips of ACC-treated F1 seedlings still staining blue (**Supplementary Fig. S5B**). This reproduces what we saw for the *EBSn:3xYPet* reporter (**Supplementary Fig. S5A**). The caveat of this experiment, again, is that *ein4* may not be fully dominant, *EIN4* may not be expressed in all cell types, and thus some cells may maintain their ethylene responsiveness, as suggested by both *3xYPet* and *GUS* versions of these reporters.

We also crossed the *EBSn:GUS* reporter with the ethylene insensitive mutant, *ein2-5*. In both control conditions and in the presence of ethylene, no GUS activity was detected in *ein2-5* seedlings in which the *EBSn:GUS* transgene was introgressed from the strongest *EBSn* line, 2.6 (**Fig. 4B-D**). We therefore conclude that the *EBSn:GUS* reporter activity in seedlings requires functional *EIN2* and is therefore ethylene-mediated. This finding further validates *EBSn:GUS* as a powerful, ethylene-specific tool that enables monitoring of plant responses to endogenous and exogenous ethylene. It is worth noting here that *ein2-5* is an EMS mutant, so we disfavor transgene silencing (which is not expected to occur in indel mutants but is relatively common in T-DNA mutant backgrounds (Schubert et al. 2004; Mlotshwa et al., 2011; Jupe et al., 2019)) as a likely alternative explanation. Unfortunately, we were not able to generate *ein2-5 EBSn:GUS* for the weaker *EBSn* line, 4.8, as the tight linkage between the transgene and *EIN2* prevented us from recombining the reporter into the *ein2* background.

Finally, we tested the rate of ethylene-triggered induction of *EBSn:GUS* in wild-type plants by evaluating the effect of shorter-term treatments with ethylene. Given the reporter’s strong basal expression level in air, we reduced GUS staining timing from overnight to 1h to avoid oversaturation. Using these conditions, we showed that the *EBSn:GUS* reporter starts responding to ethylene within 4 hours of ethylene exposure (**Supplementary Fig. S8A, B**), consistent with the ethylene induction kinetics of the genes from which the *EBSn cis*-elements were derived (**Supplementary Fig. S8C**).

### *EBSn:GUS* expression in Arabidopsis apical hooks switches off in the presence of ethylene

When working with *EBSn:GUS* lines in Arabidopsis, we made a puzzling observation that in 1-MCP and control conditions, this reporter is highly expressed in apical hooks of etiolated seedlings (**Fig. 4A**), but in ethylene (**Fig. 4A**) or ACC (**Supplementary Fig. S7**), the expression of *EBSn:GUS* in apical hooks is reduced or abolished. Thus, in the apical hooks specifically (but not in other seedling organs), low doses of ethylene appear to promote the *GUS* reporter activity, whereas high doses of the hormone appear to block it. We refer to this phenomenon as “expression pattern switching” and have observed it in all of the independent *EBSn:GUS* lines we generated (**Supplementary Fig. S7**). In fact, in stronger *GUS* lines, the shutdown of hook and cotyledon expression is often seen even at endogenous levels of ethylene, i.e. in plants grown under the control “air” conditions, with the hook expression restored only in the presence of the ethylene inhibitor 1-MCP (*EBSn:GUS* line 2.6 in **Fig. 4A**). In most extreme cases, in the presence of exogenous ethylene, the loss of GUS activity in stronger lines can sometimes expand from the hook (**Fig. 4A**) to the entire upper hypocotyl (**Fig 4B**). In contrast, we have never observed this pattern switching in any of the *3xYPet* lines, with the *EBSn:3xYPet* construct showing strong fluorescence in the apical hooks of ethylene- or ACC-treated seedlings (**Fig. 2A, B**; **Fig. 3, Supplementary Fig. S3**). Curiously, the overall *3xYPet* reporter activity pattern in ethylene-grown seedlings (**Fig. 2A**) is similar to that of moderate *GUS* lines’ staining in air (**Fig 4A**), with the predominant reporter activity observed in the cotyledons, hooks, and root tips.

We wondered what possible mechanistic reasons for the expression pattern switching in the *GUS* lines may exist. We came up with three possible explanations. First, higher levels of EIN3 accumulating in the hooks/cotyledons may result in EIN3 forming repressive transcriptional complexes that require *cis*-elements in the *GUS* open reading frame for the repression of this transgene. However, we were not able to detect any known TF-binding sites in the *GUS* construct that are not also present in the *YPet* construct. The second possibility is that excessive *GUS* (but not *3xYPet*) mRNA levels may cause local silencing in hooks/cotyledons. This explanation is unlikely, as silencing is known to spread and would be expected to cause patchy GUS activity in tissues other than the hooks, which does not appear to be the case. The third and most likely explanation is that the GUS protein, which is known to be more long-lived than standard GFP (Jefferson et al. 1987; Kavita and Burma 2008), may accumulate to very high levels in certain cells and tissues, causing protein aggregation and the inactivation of the GUS enzymatic activity. We favor the third explanation as insoluble and enzymatically inactive GUS protein aggregates have been described to form in transgenic plants under some conditions (Almoguera et al. 2002), but this is clearly not a widespread, well-described phenomenon.

To test these three possibilities experimentally, we generated yet another version of the *EBSn* reporter that expresses a *GUS-1xYPet* gene fusion (**Fig. 1B, Supplementary Table S1A**) where the two reporters are combined to encode a single chimeric protein separated by an (Gly-Gly-Gly-Gly-Ser)_2_ amino acid linker (Chen et al. 2013; Klein et al. 2014). Remarkably, the staining patterns of *GUS* and the fluorescence distribution of *1xYPet* for the fusion protein were indistinguishable from those observed in the typical individual *GUS-* and *3xYPet-*expressing lines, respectively, in both seedlings and adults (**Fig. 3**, **Fig. 4, Supplementary Fig. S9**). The only exception was that the 1xYPet fluorescence did not mark the nuclei (**Fig. 3B**) due to the lack of the nuclear localization signal in the *EBSn:GUS-1xYPet* reporter (**Fig. 1B**). These findings confirm that the apical hook switching of the GUS activity in response to ethylene is not the result of EIN3 forming the repressive transcriptional complexes in the hook nor is it a consequence of localized, hook-specific transgene silencing, as both of these scenarios would have also shut down the YPet activity in this organ (which did not happen). Therefore, we argue that the most likely reason for the expression pattern switching is the aggregation of the GUS-1xYPet fusion protein in the hook that interferes with the enzymatic activity of GUS (e.g., by limiting the accessibility of the substrate X-gluc to the fusion protein aggregate) but does not block the proper 1xYPet folding and, thus, its fluorescence (**Supplementary Fig. S10**). Regardless of the actual mechanism, our curious observation on the expression pattern switching serves as an important cautionary tale implying that the activation of *GUS* above a certain threshold may result in the loss of its enzymatic activity.

### *EBSn* is active in adult Arabidopsis plants

We next evaluated the *EBSn* promoter activity at later stages of Arabidopsis vegetative development. In plants grown in soil under standard laboratory conditions, the stronger *EBSn:GUS* reporter line, 2.6, showed a broad domain of GUS activity in young and partially expanded rosette leaves of 21-day-old plants and more localized expression at the base of fully expanded leaves (**Fig. 5A**), suggesting an active ethylene response in healthy, young rosette leaves. Consistently, GUS in the moderate *EBSn:GUS* line, 4.8, and in two *EBSn:GUS-1xYPet* fusion lines, 1 and 4, was also predominantly active in younger leaves (**Fig. 5A**). These observations suggest that younger leaf tissues may be more sensitive to (or, potentially, produce more) ethylene than older leaf tissues, an observation that we did not anticipate but that is also supported by the preferential expression of the classic *5xEBS:GUS* reporter in young leaves, although in that case GUS staining is largely confined to the leaf vasculature (**Fig. 5A**).

**Figure 5.**
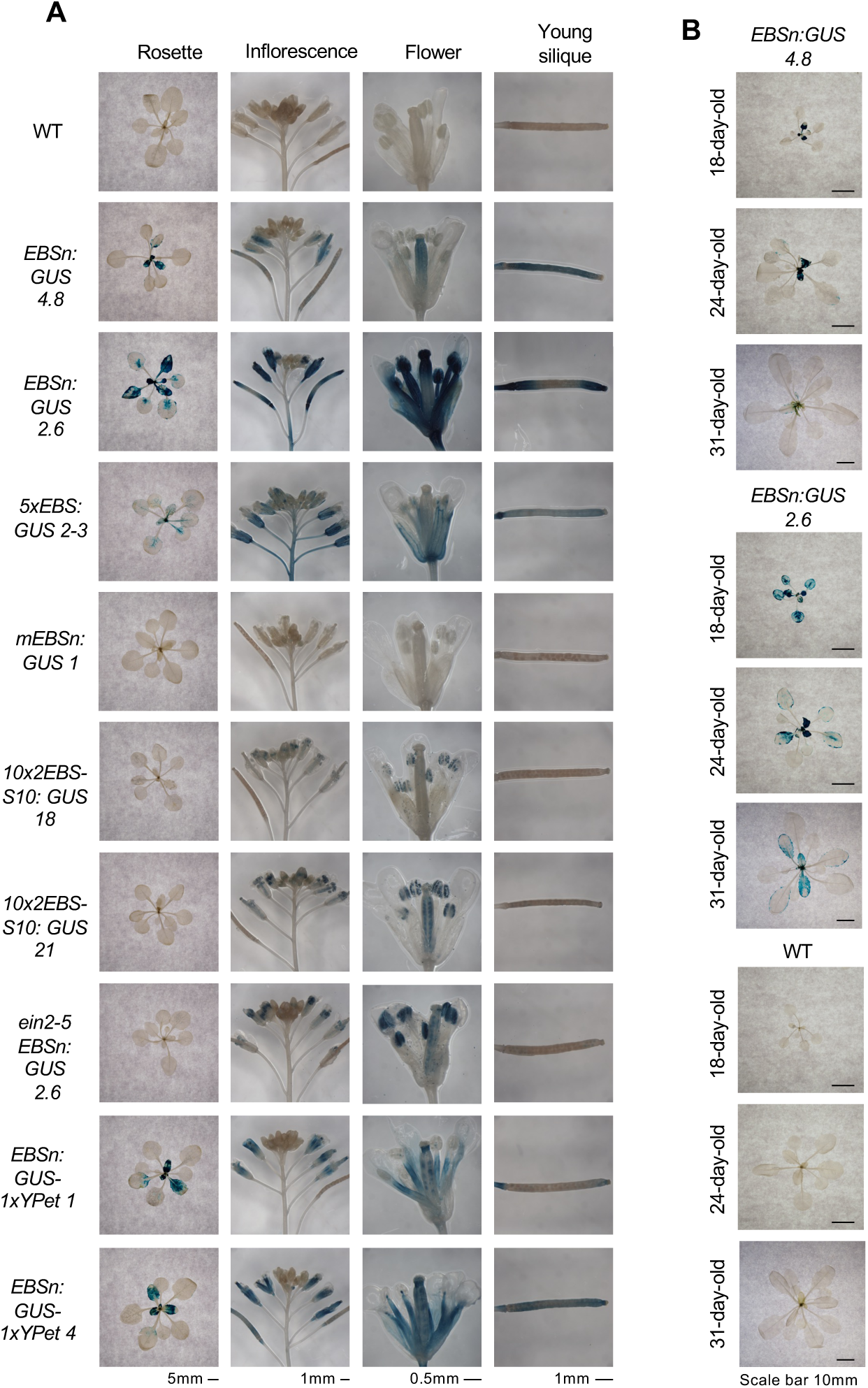
The *EBSn* promoter is active in soil-grown Arabidopsis plants. **A)** The side-by-side comparison of GUS expression patterns of *EBSn*-driven reporters and other synthetic ethylene reporters in adult *Arabidopsis* plants. First column: 21-day-old rosettes (scale bar 5mm). Second column: inflorescences of 45-day-old plants (scale bar 1 mm). Third column: close-ups of mature flowers (scale bar 0.5mm). Fourth column: close-ups of young siliques (scale bar 1 mm). **B)** Developmental regulation of the *EBSn:GUS* reporter (lines 4.8 and 2.6) in rosette leaves. GUS staining of rosettes of 18-, 24-, and 31-day-old plants of the T5 generation was compared. The staining patterns are consistent with that observed in earlier generations of *EBSn:GUS* plants. Scale bars represent 10mm. No exogenous ethylene or ACC was provided. All samples were stained for GUS overnight.

If young leaves are indeed more responsive to ethylene than older leaves, we would expect that doing GUS staining on *EBSn:GUS* rosettes at an earlier stage of development should result in more ubiquitous leaf GUS activity. Indeed, the stronger *EBSn:GUS* reporter line, 2.6, showed some staining in the entire rosette of soil-grown 18-day-old plants, but still with more prominent GUS activity detected in younger leaves (**Fig 5B**). On the other hand, the weaker of the two *EBSn:GUS* lines, 4.8, displayed GUS activity exclusively in young leaves of 18-day-old plants, confirming that young leaves may be more responsive to ethylene than older leaves (**Fig 5B**). As seen before with older rosettes of *EBSn:GUS*, by 24 days of age, both lines restricted their GUS activity to younger leaves only, and such reporter expression pattern persisted in 31-day-old rosettes (**Fig 5B**).

We also examined the basal levels of YPet reporter activity in 45-day-old rosette leaves of soil-grown *EBSn:3xYPet* (lines A and C) and *EBSn:GUS-1xYPet* (lines 1 and 4) plants, but saw no prominent leaf fluorescence in any of the four lines under standard growth conditions, apart from some activity at the site of the cut in the petiole (**Supplementary Fig. S11A**). However, as anticipated, upon 24-hour-long exposure of detached leaves to exogenous ACC, the two *3xYPet* lines showed increased leaf fluorescence. In contrast, the *1xYPet* signal in the two *GUS-1xYPet* lines was not easily detectable in the leaf blade and was primarily restricted to the damaged petiole (**Supplementary Fig. S11A**).

For GUS reporters, the histochemical staining in ACC-treated, detached, 45-day-old rosette leaves of the moderate *EBSn:GUS* line, 4.8, and of the two *EBSn:GUS-1xYPet* fusion lines, 1 and 4, revealed upregulated levels and expanded domains of GUS activity relative to their mock-treated controls, consistent with the notion that the ACC treatment activated ethylene responses (**Supplementary Fig. S11B**). In contrast, the ACC treatment did not broaden the GUS expression domain in the leaves of the stronger *EBSn:GUS* line, 2.6, suggesting that in this line the maximum level of *GUS* activity was already achieved in control conditions. Consistent with these observations, damaging 45-day-old rosette leaves by smashing them with forceps, a treatment that is expected to lead to a boost in endogenous ethylene production and signaling in wounded tissues (Di Fino et al 2025), resulted in an increase in the *EBSn:GUS* activity only in the moderate *EBSn:GUS* line, 4.8, but not in the saturated stronger line, 2.6 (**Supplementary Fig. S11B**). The lack of increased expression in damaged leaves of the *EBSn:GUS* 2.6 line supports the notion that in that line the reporter activity is already maximized at lower endogenous levels of ethylene. Given the characteristic patchy (strong at the base, absent at the tip) domain of GUS activity in this line and our inability to trigger enhanced or expanded GUS staining with ACC treatment or deliberate tissue damage, we argue that the aforementioned GUS inactivation, like that observed in apical hooks of the *EBSn:GUS* line 2.6, may also be happening in leaves, possibly due to GUS aggregation (**Supplementary Fig. S10**) or reporter silencing.

To determine whether the activity of *EBSn:GUS* and *EBSn:3xYPet* in rosette leaves, like that in seedlings, is ethylene-dependent, we examined the expression of these reporters in the *ein2-5* background. No basal GUS or 3xYPet reporter activity was observed in the rosette leaves of soil-grown 21- and 45-day-old plants harboring either of the *EBSn* transgenes introgressed via crossing into *ein2,* and the reporters were not induced upon ACC treatment or leaf mechanical damage (**Fig. 5A, Supplementary Fig. S11**). These findings confirm the indispensable role of *EIN2* in ethylene signaling and demonstrate the requirement for functional *EIN2* for the transcriptional response of the *EBSn*-driven reporters to ethylene. Furthermore, we tested the leaves of the *mEBSn:GUS* mutant reporter, and as expected, saw no reporter activity in these plants that carry a defective version of the *EBSn* reporter in which all EIN3/EIL binding sites in the synthetic promoter are inactivated (**Fig. 5A**), consistent with the notion that EIN3/EIL binding to the promoter is necessary for the reporter activity.

Finally, we examined the inflorescences of wild-type *EBSn:GUS* reporter lines grown under standard laboratory conditions in soil and discovered that the stronger line, 2.6, had high levels of expression in post-anthesis flowers, anther-specific expression in pre-anthesis flowers, and no prominent expression in young flower buds (**Fig. 5A**), suggesting that only some flower tissues can support expression of the *EBSn:GUS* reporter under endogenous ethylene concentrations. In the weaker line, 4.8, the reporter expression was even more restricted and prominent only in the style of post-anthesis flowers (**Fig. 5A**), confirming that *EBSn:GUS* is preferentially triggered in a subset of flower tissues, which is indicative of a refined spatio-temporal activation of the ethylene pathway during normal developmental processes. In comparison, the sepal-enriched pattern of expression of *5xEBS:GUS* and anther-specific activity of *10x2EBS-S10:GUS* both stood in sharp contrast to that of *EBSn:GUS.* Such discrepancies are, in our opinion, not unexpected and are likely reflective of the differential activity of alternative EIN3/EIL-containing transcriptional complexes forming in different flower tissues and having specific binding preferences for certain promoter architectures, as already noted above for seedlings. In contrast, what was very surprising for us to discover is that in the *ein2*-*5* background, *EBSn:GUS* shows prominent residual expression specifically in the anthers/pollen of post-anthesis flowers, a pattern similar to that observed in the flowers of two different *10x2EBS-S10:GUS* in the wild-type background (**Fig. 5A**). It is currently unclear whether the EIN2-independent expression of *EBSn:GUS* in the anthers is triggered by ethylene or by another environmental or developmental signal. However, given that the *mEBSn:GUS* negative control is devoid of any activity in all seedling and adult tissues tested, including flowers (**Fig. 5A**), we conclude that the activity of the *EBSn* promoter is fully dependent on EIN3/EILs binding and does not stem from the basal activity of the *35S* core promoter alone.

### *EBSn* is functional in tomato

To assess the utility of the *EBSn* reporters in species beyond Arabidopsis, we transformed the *GUS* and *3xYPet* versions of the constructs (**Supplementary Table S1B**) into tomato and generated five *GUS* lines and ten *3xYPet* lines (**Supplementary Fig. S12**). All screened *EBSn:GUS* lines displayed some degree of transgene silencing either for the reporter, the kanamycin resistance, or both (**Supplementary Fig. S13**). Therefore, we selected *EBSn:GUS* and *EBSn:3xYPet* lines for further characterization in T2 based on the consistency of their reporter expression patterns and levels across generations. In parallel, to prevent potential silencing of the transgene, we backcrossed selected lines to wild type (M82) (**Supplementary Fig. S14**) and isolated stable single-insertion events (**Supplemental Table S2**).

Examination of the expression patterns of the *EBSn:GUS* reporter in etiolated tomato seedlings in control conditions revealed similar GUS staining patterns (**Fig. 6A**) to that of the respective Arabidopsis lines (**Fig. 4**), with the highest basal reporter activity detected in the apical hooks and root tips and further expansion of the domain of GUS expression observed upon exposure of seedlings to exogenous ACC. Likewise, easily detectable *EBSn:GUS* reporter activity was observed in the developing tomato leaves of three-week-old light-grown plants at endogenous levels of ethylene (**Fig. 6B, C**), which is consistent with the aforementioned high levels of basal GUS activity seen in young rosette leaves of *EBSn:GUS* lines in Arabidopsis (**Fig. 5**). Not surprisingly, a robust upregulation of GUS staining was seen in tomato leaves exposed to 50 μM ACC for 24 hours (**Fig. 6C)**, which was recapitulated in the equivalent *EBSn:3xYPet* lines of tomato (**Fig. 6D**), consistent with the results observed in Arabidopsis rosette leaves **(Supplementary Fig. S11B**). These findings suggest that both the *GUS* and *3xYPet* reporter versions are functional in tomato even though the *EBSn* promoter was generated from Arabidopsis-derived ethylene-responsive *cis*-elements. Noteworthy, we found that in tomato leaves stomata guard cells show the highest level of 3xYPet fluorescence in response to exogenously provided ACC (**Supplementary Fig. S12 and S13**), but it remains to be determined if these cells are *de facto* more sensitive to ethylene than the surrounding epidermal pavement cells or simply uptake more ACC from the media.

**Figure 6.**
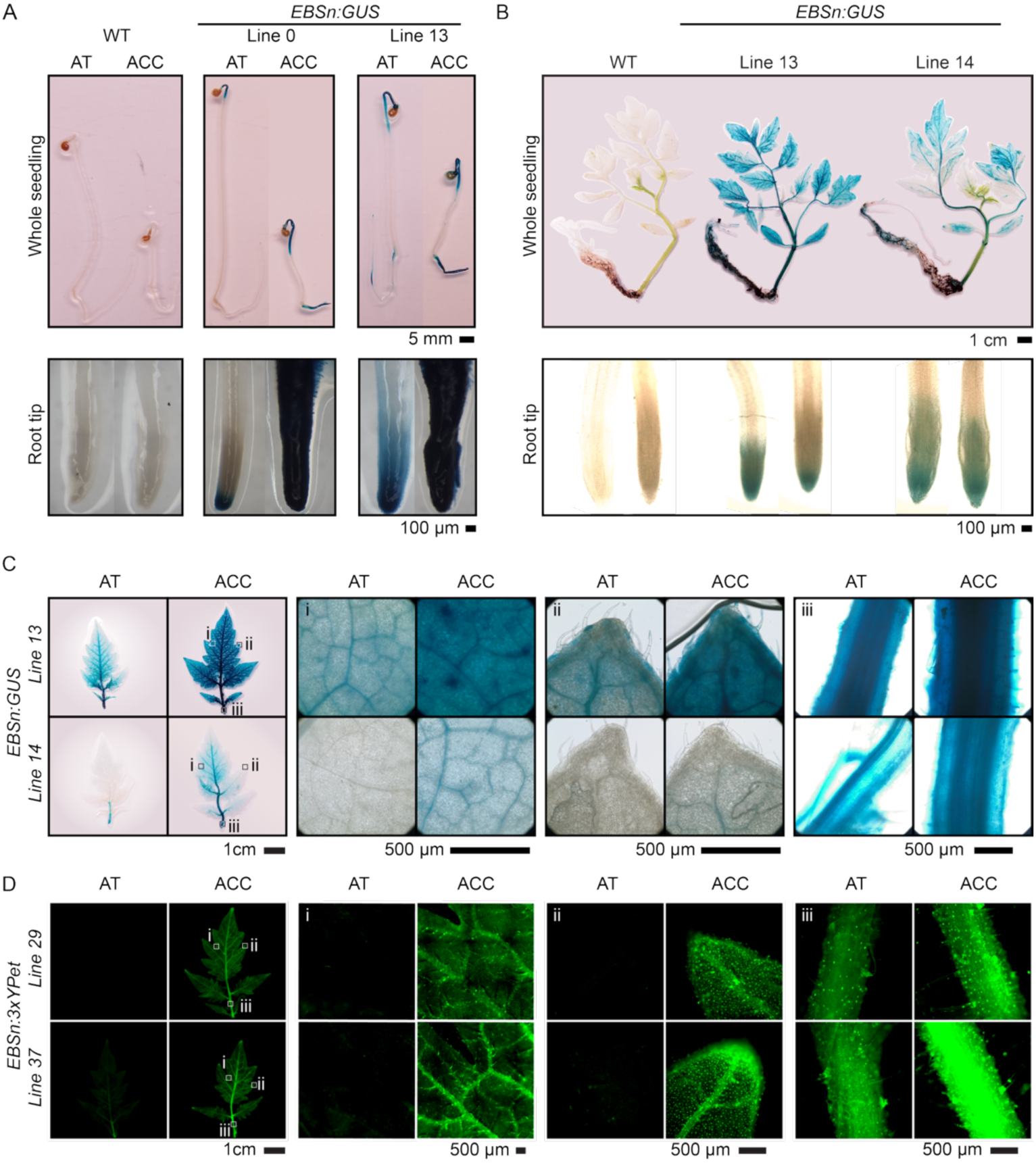
The *EBSn:GUS* reporter is expressed in tomato and inducible by exogenous ACC. **A)** The *EBSn* promoter is active in three-day-old etiolated T2 tomato seedlings and strongly inducible by 1 µM ACC (equivalent to 0.1 µM ACC in Arabidopsis based on ACC/control relative root length). Two independent *EBSn:GUS* transgenic lines (0 and 13) germinated for three days in the dark in magenta boxes containing either plain AT or AT supplemented with ACC is displayed. GUS staining was performed overnight. **B)** *EBSn:GUS* is active in leaves and roots of three-week-old T2 tomato plants (lines 13 and 14) grown under long day conditions in soil. Seeds were germinated in AT media supplemented with 100µg/mL kanamycin (except for WT germinated in plain AT media) in the dark for three days and transferred to continuous light for an additional week. After a total of 10 days, seedlings were transferred to soil and grown at 22 °C in long day conditions (16-h light:8-h darkness) for four more days. GUS staining was performed overnight. **C,D)** The *EBSn* activity is strongly induced in the vasculature and rachis of detached three-week-old leaves of T2 tomato plants (lines 29 and 37) upon 24-hour exposure to 50 µM ACC. Leaves were detached, and the abaxial side of the leaf was imbibed for 24 hours in plain 0.6% (w/v) bactoagar or bactoagar supplemented with 1 µM ACC. Images of GUS staining (panel C) or 3xYPet fluorescence (panel D) are displayed. WT = *Solanum lycopersicum* M82.

When analyzing the GUS staining in roots of three-week-old soil-grown *EBSn:GUS* tomato seedlings, we found that the level of GUS activity under ambient conditions often varied from one tomato root to another (**Fig. 6B**). We speculate that this may be the intrinsic property of different lateral root orders (e.g., secondary versus tertiary) or, potentially, the consequence of the structural heterogeneity of the soil, with different levels of local soil compactness affecting the accumulation of ethylene (Pandey et al. 2021). In fruits, consistent with tomato being a climacteric species that produces high levels of ethylene at the onset of ripening (Alexander and Grierson 2002; Quinet et al. 2019), we observed strong *EBSn:GUS* and *EBSn:3xYPet* activity in ripe or ripening fruits (**Fig. 7A**), further validating the utility of *EBSn* in visualizing responses to both endogenous and exogenous ethylene in different organs of this species.

**Figure 7.**
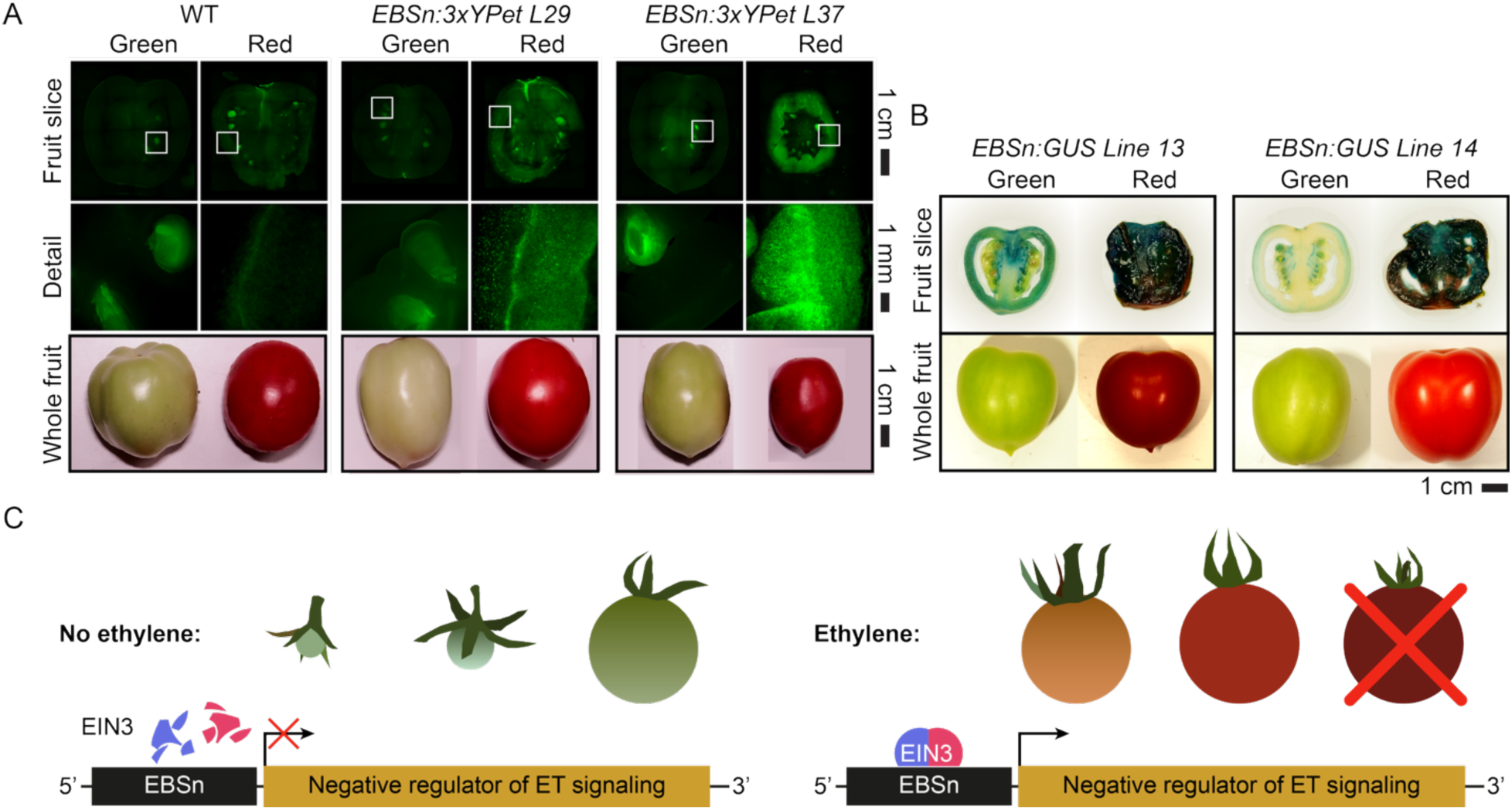
The *EBSn* activity in tomato fruits is enhanced upon ripening. **A)** YPet fluorescence in T2 *EBSn:3xYPet* tomato lines 29 and 37 is weak in green fruits and strongly induced in red fruits. **B)** GUS staining in T2 *EBSn:GUS* tomato lines 13 and 14 is weak in green fruits and prominently induced in red fruits. Fruit slices were fixed and stained for GUS for one hour. **C)** Potential future application of the *EBSn* synthetic promoter. By expressing a negative regulator of ethylene signaling under the control of *EBSn*, it should be possible to counteract the effects of ripening-induced ethylene and avoid over-ripening. In developing green fruits, the ethylene production is low, the EIN3 TF is destabilized, and the expression of a negative regulator such as *EBF1/2* (or, alternatively, of an antisense version of a positive regulator such as *EIN2*) is off, and fruits develop normally. In maturing fruits, ethylene production and signaling are triggered at the onset of fruit ripening, thus stabilizing EIN3/EILs and activating the expression of the negative regulator (or the antisense version of the positive regulator), inhibiting further ethylene signaling, and preventing fruit over-ripening. As a result, transgenic fruits should develop normally until the boost in ethylene production at the onset of ripening, at which point ethylene would self-inhibit further ethylene signaling, delaying fruit decay. WT = *Solanum lycopersicum* M82.

### Final conclusions and outlook

Herein, we report the generation and initial characterization of a synthetic ethylene-inducible promoter, *EBSn*, in Arabidopsis and tomato. The promoter is sensitive and robust in its response to ethylene, making it a great tool for monitoring plant responses to endogenous or exogenous ethylene. Our reporter characterization indicates that the *EBSn* promoter activation requires EIN3 binding since point mutations in the EIN3 target site in *mEBSn* abolish promoter function. We also conclude that in dark-grown seedlings and soil-grown rosette-stage adult plants, this promoter’s activity is fully dependent on *EIN2,* implying that the high basal activity of the *GUS* version of the *EBSn* reporter is triggered by endogenous ethylene rather than another developmental or environmental stimulus. The absolute dependence of this promoter activity on functional *EIN2* is however broken in the pollen, as in post-anthesis flowers of soil-grown plants, there is restricted but prominent *EBSn:GUS* activity in the *ein2-5* mutant pollen. *ein2-5* is a well-characterized genetic background that was shown to be fully ethylene insensitive in most (Alonso et al. 1999; Althiab-Almasaud et al. 2021; Li et al. 2022) but not all (Kim et al., 2013) genetic studies. Therefore, it remains to be determined whether the residual GUS activity of *EBSn:GUS* in the *ein2-5* pollen is ethylene-mediated, but the promoter activity is disrupted by point mutations in the EIN3 binding sites in *mEBSn:GUS* lines.

The *3xYPet, GUS* and *GUS-1xYPet* versions of the *EBSn* reporter described in this work appear to be stable across generations, and we envision that these can serve as convenient tools to evaluate ethylene responsiveness of wild-type plants and mutants alike (as we did herein with *ein4* and *ein2*), or to study the role of ethylene in plant development (e.g., in seed germination, organ abscission, or fruit ripening), as well as in responses to pathogens, endophytes, abiotic stress, wounding, parasitic plants, insect damage, etc. Although the *EBSn* promoter *cis*-elements were sourced from Arabidopsis genes, the high degree of conservation of TF-binding sites across different plant species (Mao et al. 2022) makes us optimistic about the utility of this tool in many plants, as supported by our *EBSn* reporter analysis in tomato.

In the future, besides its utility in biosensing, the *EBSn* promoter has the potential to aid in reducing food waste caused by produce senescence and fruit overripening. One can envision, for example, engineering a fruit tissue-specific negative feedback loop that delays ripening or senescence-induced processes in response to endogenous or exogenous ethylene. By driving the expression of negative regulators of ethylene signaling (such as *EBF1/2* or dominant versions of any of the ethylene receptor genes) or antisense constructs of positive regulators (such as *EIN2* or *EIN3*) under the control of the *EBSn* promoter, one should be able to inhibit ethylene signaling in response to ethylene accumulation, creating a negative feedback regulatory loop that effectively slows down the undesired processes of overripening and senescence (**Fig. 7B**). The trick would be to target the ethylene pathway autoinhibition to specific tissues and developmental stages (e.g., mature fruits), which should be possible with the implementation of basic genetic logic gates in plants (Zhao et al. 2021). We are optimistic about the potential of synthetic biology to drive biotech innovation and foresee the implementation of a wide array of multi-gene circuits in plants in the near future.

## Materials and methods

### Design of the *EBSn* and *mEBSn* synthetic proximodistal promoters

We set out to build a novel synthetic ethylene-responsive promoter with enhanced sensitivity and broad domain of activation. To generate *EBSn*, the following design pipeline was implemented. 21bp-long natural DNA elements from the Arabidopsis genome were extracted from EIN3 ChIP-seq data (Chang et al. 2013). These sequences consist of an 11bp-long natural *2EBS(−1)* element flanked by 5bp-long spacers naturally present in the genome upstream and downstream of the 11bp core (**Fig. 1A, Supplementary Fig. S2A**). The following two criteria were applied as filters to retrieve the best *2EBS(−1)* elements: (1) the sequences are located within the EIN3 ChIP-seq peaks; and (2) those *2EBS(−1)* sequences fall in the promoters of genes strongly inducible by ethylene (Chang et al. 2013). 18 of these 21bp sequences derived from 14 best genes (**Supplementary Table S3, Supplementary Fig. S8C**) that pass these two criteria were randomly stacked together in groups of 10, potentially unintentionally generating other TF-binding sites at the junctions. To identify the possible TF-binding sites created at the junction sequences, we bioinformatically screened the resulting *10xEBS* sequences for the presence of EIN3-unrelated TF-binding motifs using position weight matrix models (Wasserman and Sandelin 2004). The *10xEBS* combinations that lead to the highest ratio between the frequency of the predicted EIN3-binding motifs and the frequency of the predicted undesired TF-binding sites (O’Malley et al. 2016; Tsukanov et al. 2022) were kept, and key nucleotides within the latter sequences were replaced to abolish undesired TF binding (**Supplementary Fig. S2A, Supplementary Table S4**). The source code implementing the bioinformatics approach used to generate *EBSn* and *mEBSn* sequences is available at https://github.com/parthian-sterlet/gsga.

The resulting sequence, *EBSn*, is a tandem of 10 semi-synthetic, buffered *2EBS(−1)* elements spaced in a manner that places EIN3 dimers on the same side of the DNA double helix, with each 21bp imperfect repeat taking exactly two helical turns (**Fig. 1A, Supplementary Fig. S2A**). In parallel with designing *EBSn,* its mutant version, *mEBSn*, was developed in which one conserved nucleotide in each half-site of the *2EBS(−1)* core element was intentionally mutated with the goal to abolish EIN3 binding, thus creating an EIN3-blind negative control (**Fig. 1A, Supplementary Fig. S2B**).

### Plasmid construction and transgenic line generation

The promoter sequences for *EBSn* and *mEBSn* (**Supplementary Fig. S2A, B**) were domesticated *in silico* for their compatibility with the GoldenBraid (GB) cloning system (Sarrion-Perdigones et al. 2013; https://gbcloning.upv.es/), commercially synthesized as a gBlock (IDT Technology, https://eu.idtdna.com/pages), and cloned into the GB entry level vector, *pUPD2* (**GB0307**) (**Supplementary Table S1**). The primers used to amplify the *de novo*-synthetized *EBSn* and *mEBSn* promoters from the gBlock (**Supplementary Table S4**) harbored a 5’GB overhang in the forward oligo (which included a BsmBI restriction site, a *pUPD2*-compatible grammar code, and a 5’-[A1] GB grammar code, followed by the sequence of the first *EBS* repeat, intact or mutated) and a 3’ GB overhang in the reverse oligo (which contained the BsmBI restriction site, the *pUPD2* grammar, and the 3’-[A2] GB grammar, followed by the sequence of the last *EBS* repeat, intact or mutated). In parallel, two additional *EBSn-mini35S\** and *mEBSn-mini35S\** sequences (i.e., functional and mutated *EBSn* versions fused with the *mini35S\** core promoter variant (**Supplementary Fig. 1C**) used in the *5xEBS* original reporter (Stepanova 2001; Stepanova et al. 2007)) (**Fig. 1B**) were also synthesized as part of the same gBlock. To amplify the latter two sequences from the gBlock, the aforementioned forward oligos were used in combination with a new reverse oligo with similar 3’ end but followed by the *mini35S*\* sequence instead of an *EBSn* or an *mEBSn* repeat. The sequences for all four promoters synthesized as parts of the same gBlock and corresponding primers can be found in **Supplementary Table S4**. The resulting *pUPD2_EBSn*, *pUPD2_mEBSn*, *pUPD2_EBSn:mini35S*,* and *pUPD2_mEBSn:mini35S\** [A1-A2] entry clones were assembled into transcriptional reporters in the alpha-level destination vector *pDGB3alpha1* together with the following DNA parts: *pUPD2_mini35S* (−46, UTR) [A3-B1] as core promoter (**Supplementary Fig. 1C**), *pUPD2_3xSV40-NLS* [B2] as nuclear localization signal, *pUPD2_3xYPet* or *pUPD2_intronless*-*GUS* [B3-B5] (**GB0208**) as CDS, and *pUPD2_term35S* [C1] (**GB0036**) as terminator (Franck et al. 1980, Odell et al. 1985, Jefferson et al. 1987, Hicks and Raikhel 1993, Nguyen and Daugherty 2005, Eudes et al. 2008, Zhou et al. 2011, Brumos et al. 2020; Fernandez-Moreno et al. 2024) following the previously described procedure (Fernandez-Moreno et al. 2024). The resulting fluorescent or histochemical transcriptional reporters were tested for activity by tobacco leaf agroinfiltration and assembled in an omega-level destination vector *pDGB3omega1* or *pDGB3omega2* with the selectable marker *pDGB3alpha2_tNOS:BASTA:pNOS* (cloned into alpha level from the *pUPD* entry clone **GB0023**; Fernandez-Moreno et al. 2024) for Arabidopsis transformation or with *pDGB1alpha2_pNOS:KANAMYCIN:tNOS* (**GB0184**) for tomato transformation (see omega-Resistance module in **Supplementary Table S1**).

To make the double *EBSn:GUS-YPet* reporter, *pUPD2_EBSn* [A1-A2], *pUPD2_mini35S* (−46, UTR) [A3-B1] and *pUPD2_term35S* (**GB0036**) [C1] were used in combination with two new DNA parts: *pUPD2_intronless-stopless-GUS* [B2] and *pUPD2_(GGGS)x2-1xYPet* [B3-B5] as CDS (no nuclear localization signal was used for this double reporter) (see entry clones and alpha-level reporters in **Supplementary Table S1**). The *pUPD2_intronless-GUS* [B3-B5] (**GB0208**) was used as a template to amplify the *GUS* sequence using “GUS_B2F: GCGCCGTCTCGCTCGCCATGTTACGTCCTGTAGAAACC” and “GUS_B2R: GCGCCGTCTCGCTCACATTTGTTTGCCTCCCTGCTGCG”. The reverse oligo was designed to exclude the final stop codon (TGA). The *pUPD2_(GGGGS)x2-1xYPet* [B3-B5] clone was created using the GGGGS-YPET_B3F: GCGCCGTCTCGCTCG<u>AATG</u>GGAGGTGGAGGTTCGGGTGG and YPET_B5R: GCGCCGTCTCGCTCA<u>AAGCT</u>CACTTATACAATTCATTCATTCCTT forward and reverse primers, respectively. An existing lab clone which contains an in-frame Gly-Gly-Gly-Gly-Ser (1xGGGGS) linker upstream of a *1xYPet* fluorescent protein gene was used as a PCR template. The forward oligo added an additional 1xGGGGS linker during the amplification, resulting in a final 2xGGGGS-*1xYPet* clone. Both sets of oligos harbor the GB3.0 entry-level overhangs at their 5’ and 3’ flanks, including the BsmBI restriction site, the 4nt grammar code compatible with the *pUPD2* entry vector, and the 4nt B2 and B3-B5 grammar codes for the *GUS* and *1xYpet* clones, respectively (see underlined in GGGGS-YPET B3F and YPET B5R oligonucleotide sequences). The different entry clones, *pUPD2_EBSn* [A1-A2], *pUPD2_mini35S* (−46, UTR) [A3-B1], *pUPD2_intronless-stopless-GUS* [B2], *pUPD2_(GGGGS)_2_-1xYPet* [B3-B5], and *pUPD2_term35S* [C1], were next assembled into a double transcriptional reporter (**Fig. 1B**) in the alpha-level destination vector *pDGB3alpha1*, and then into the omega-level destination vector *pDGB3omega1* in combination with the *pDGB3alpha2_tNOS:BASTA:pNOS* clone (Fernandez-Moreno et al. 2024) to create the final expression module (**Supplementary Table S1A**).

The resulting modules were transformed into *Agrobacterium tumefaciens* strain GV3101 via electroporation (Alonso and Stepanova 2015). Positive clones were transformed into Arabidopsis Columbia-0 plants via floral dip (Clough and Bent 1998) or into the tomato M82 cultivar via tissue culture (Krasnyanski et al. 2001; Van Eck et al. 2019).

Transgenic T1 Arabidopsis lines were selected in AT plates (1xMS (4.33 g L^-1^ Murashige & Skoog media (PhytoTechnology Lab)), pH 6, 1% sucrose, 6 g L^-1^ Bactoagar) supplemented with 10-20 ug/ml phosphinotricin (the active ingredient in Basta). Homozygotes were identified in T3 and carried to the T4 generation. Plants were propagated in Sun Gro Professional Mix soil under 16h/8h light/dark cycle. For an extended description of these procedures, see Fernandez-Moreno et al. (2024). All in all, four types of Arabidopsis transcriptional reporter lines were generated: one fluorescent reporter (*EBSn:mini35S:3xNLS:3xYPet:term35S,* elsewhere referred to as *EBSn:3xYPet* (**Fig. 1 B1**)), two histochemical reporters (*EBSn:mini35S*:3xNLS:GUS:term35S,* referred to as *EBSn:GUS* (**Fig. 1 B2**), and its mutant version *mEBSn:mini35S*:3xNLS:GUS:term35S,* referred to as *mEBSn:GUS* (**Fig. 1 B5**)), and one combined fluorescent-histochemical reporter fusion (*EBSn:mini35S:GUS-1xYPet:term35S,* referred to as *EBSn:GUS-1xYPet* (**Fig. 1 B6**)). Arabidopsis *5xEBS:GUS* (Stepanova 2001; Stepanova et al. 2007) (**Fig. 1 B3)** and *10x2EBS-S10:GUS* lines (Fernandez-Moreno et al. 2024) (**Fig. 1 B4**) were reported previously.

Transgenic T0 tomato plants that formed well-developed roots in Tomato Rooting (TR) medium (4.4 g/L MS salts + Gamborg’s vitamins, 30 g/L sucrose, and 8 g/L agar, pH 5.6) supplemented with 300 mg/L timentin and 100 mg/L kanamycin were transferred individually into small plastic pots prefilled with sterile Sun Gro Professional Mix soil, covered with clear plastic for a week, and grown in a growth chamber under a 16-hour light photoperiod (20 W cool-white fluorescent light, 30 μmol m-2 s-1) at 25°C until plants reached the height of ∼20-25 cm. At that point, plants were transferred to 7.57 L-pots containing Sun Gro Professional Mix soil, one plant per pot, moved to the greenhouse, and grown under ambient light or supplemented light during winter months to guarantee 16-hour light cycles at 18°C target temperature, with highs ∼32°C in the summer and lows ∼15°C in the winter. Two types of tomato lines were produced, those that harbor *EBSn:mini35S:3xNLS:3xYPet:term35S,* aka *EBSn:3xYPet* (**Fig. 1 B1**), and those that carry *EBSn:mini35S*:3xNLS:GUS:term35S,* aka *EBSn:GUS* (**Fig. 1 B2**).

### Hormone inducibility and wounding assays in transgenic Arabidopsis

Arabidopsis seeds from the T2 to T5 generations of the *3xYPet, GUS,* and *GUS-1xYPet* fusion reporter lines were surface-sterilized with bleach following previously described protocols (Brumos et al. 2020). For inducibility assays using ACC, the sterilized seeds were plated on control AT plates and AT plates supplemented with the indicated concentrations of ACC (PhytoTechnology Lab), typically 10 μM. For assays involving 10 ppm ethylene gas (Airgas), hydrocarbon-free air, or 200 ppm 1-MCP (AgroFresh), seeds were plated on plain AT plates. All plated seeds were stratified at 4 °C in the dark by placing stacks of aluminum-foil wrapped plates in the cold for two to five days, then exposed to light for 1-2 hours at room temperature to promote germination and subjected to their respective treatments at 22 °C in the dark for three days (72h), unless otherwise noted. For a 1-MCP treatment, a single 1.25 g sachet of SmartFresh InBox powder (AgroFresh, 0.014% 1-MCP) was combined with ∼2ml of water in an airtight 1L container with plates. For gas treatments, plates were placed into individual airtight 1L containers equipped with an inlet and an outlet, and flow-through hydrocarbon-free or ethylene-supplemented (10 ppm) air was applied for an indicated amount of time (typically 72h, unless indicated otherwise).

Leaf explants from 45-day-old, soil-grown adult Arabidopsis plants were harvested and either deliberately damaged with forceps by gently squashing the leaf blade in several locations of the leaf blade or left intact, placed on the surface of 8 g/L agar plates with or without 10 μM ACC, and incubated for 24 hours to study either GUS activity as previously described (Brumos et al., 2020) or YPet fluorescence using a Leica Thunder Imager M205FA equipped with DFC9000 sCMOS camera (fluorescence).

### Arabidopsis GUS staining and imaging

Three-day-old, etiolated Arabidopsis seedlings from *EBSn:GUS*, *mEBSn:GUS*, *EBSn:GUS-1xYPet, 10x2EBS-S10:GUS* (Fernandez-Moreno et al. 2024), and *5xEBS:GUS* (Stepanova et al. 2007) reporter lines (T2 to T4 generations) exposed to different treatments were fixed in ice-cold 90% acetone and stained for GUS for an indicated amount of time (1h to overnight) at 37 °C as described (Stepanova et al. 2005). Stained seedlings were either directly imaged or first optically cleared using a freshly prepared ClearSee solution (Kurihara et al., 2015) for 7 days before imaging (Brumos et al. 2020). GUS-stained seedlings were either mounted on glass slides and imaged using a 5.0 RTV digital camera (Q Imaging, Surrey, BC, 904 Canada) under a Zeiss AxioSkop2 Plus microscope with Nomarski optics (Brumos, Zhao et al., 2019) (**Supplementary Fig. S8B**) or displayed on agar plates and imaged using a 5.0 RTV digital camera (Q Imaging, Surrey, BC, 904 Canada) under a Stereo Zoom Microscope (Leica mZ125) (**Supplementary Fig. S8A**).

Rosettes of two- to four-week-old and inflorescences of 45-day-old soil-grown Arabidopsis reporter lines *EBSn:GUS*, *mEBSn:GUS*, *EBSn:GUS-1xYPet, 10x2EBS-S10:GUS* (Fernandez-Moreno et al. 2024), and *5xEBS:GUS* (Stepanova et al. 2007) were fixed for one week in 90% acetone to remove chlorophyll and stained for GUS as described above for seedlings. Rosettes were floated in a 70% ethanol solution and imaged using a digital rangefinder-style mirrorless camera (Olympus PEN E-PL6) (**Fig. 5**). Inflorescences, flowers, and young siliques were displayed on agar plates and imaged using a 5.0 RTV digital camera (Q Imaging, Surrey, BC, 904 Canada) under a Stereo Zoom Microscope (Leica mZ125) (**Fig. 5A**).

### YPet fluorescence imaging of Arabidopsis seedlings

Three-day-old, etiolated seedlings from *EBSn:3xYPet* and *EBSn:GUS-1xYPet* (T2 to T4 generations) fluorescent reporter lines germinated under different treatments were mounted on glass slides in water and imaged using a DFC365 FX camera and a Zeiss Axioplan microscope (**Fig. 2A-B** and **Supplementary Fig. S3, S5A, S12B**), a Leica Thunder Imager M205FA equipped with DFC9000 sCMOS camera (**Supplementary Fig. S4, S6, S9, and S11A**), or a Zeiss LSM880 confocal microscope at 1 aerial unit (AU) (excitation/emission wavelengths of 488 nm/492–570 nm) (**Fig. 2C-D and 3B**). ImageJ v1.52, Fiji 2015 (**Figures 2A-B and 3B; Supplementary Fig. S3A-B**) and LEICA LAS X software (**Supplementary Figs. S4, S6, S9, and S11A**) were used for image processing.

### Tomato growth conditions

Tomato seeds were surface-sterilized for three minutes in 70% ethanol followed by 7 minutes in 30% commercial bleach (1.8% sodium hypochlorite final concentration), washed 5 or more times with sterile water, and stratified for 3-5 days at 4°C. Seeds were next placed with forceps on vertical AT plates (4.4 g/L MS salts, 10 g/L sucrose, pH 6, 10 g/L bacto-agar) supplemented with 100 mg/L kanamycin. Plates with seeds were then positioned vertically under continuous LED light, 70-100 µmol m-2 s-1 (2x 6000K Kihung T8 LED integrated fixture 40W + 1x FULL SPECTRUM Monios-L LED grow light full spectrum 60W) and incubated for seven days.

Kanamycin-resistant plants were identified for their longer roots and healthier leaves, transferred into small plastic pots (0.5 L) filled with sterile soil (1:1 ratio of Sun Gro professional growing mix and Jolly Gardener Pro-Line C/B growing mix), and kept under clear plastic covers for at least a week in the growth chamber under 100-120 µmol m-2 s-1 LED light in a 16-h light/8-h dark photoperiod under the aforementioned light fixture. Once plants reached a height of approximately 20-25 cm, they were transplanted to 7.57 L black plastic pots filled with Sun Gro Professional Growing Mix and hand-watered every two-three days as needed to prevent wilting. Plants were fertilized biweekly with Bloom City CAL-MAG 2-0-0 to prevent tomato blossom end rot caused by calcium deficiency.

### Tomato treatment and imaging

For seedling imaging, T2 tomato seeds were surface-sterilized as described above, placed in transparent magenta boxes containing control AT medium or AT supplemented with 1 µM ACC, and exposed for two hours to ambient light at room temperature to promote germination. After this, the boxes with seeds were either transferred to the dark for three days or placed under continuous LED light for five days. *EBSn:GUS* and *EBSn:3xYPet* leaf explants were placed on the surface of 8 g/L agar plates with or without 50 μM ACC and incubated for 24 hours. Next, *EBSn:GUS* seedlings, leaf explants, whole three-week-old plants, and fruits divided in ∼2 mm-wide slices were fixed in 90% (v/v) acetone, stained for GUS as described (Brumos et al. 2020), and imaged using an OLYMPUS PEN Lite E-PL6 digital camera (**Fig. 6A-B** top panels, **Fig. 6C** leftmost panel, **Fig. 7B**; **Supplementary Fig. S12A and S13A**) or using a Diagnostic Instruments Spot Insight 4 14.2 color mosaic camera coupled to a Zeiss Axioplan microscope (**Fig. 6A-B** bottom panels, **Fig. 6C** detail panels). *EBSn:3xYPet* leaves and fruit sections were imaged live without prior fixation using a Leica Thunder Imager M205FA equipped with DFC9000 sCMOS camera (fluorescence, objective 1X, zoom 2X, exposure 3 seconds) (**Fig. 6D** and **7A**; **Supplementary Fig. S12B**), and a Zeiss LSM880 confocal microscope at 1 aerial unit (AU) (excitation/emission wavelengths of 488 nm/499–562 nm) (**Supplementary Fig. S12C and S13B**). ImageJ v1.52, Fiji 2015 (**Supplementary Fig. S12C and S13B**) and LEICA LAS X software (**Fig. 6D and Fig. 7A; Supplementary Fig. S12B**) were used for image processing.

## Acknowledgements

A.E.Y. is grateful for her fellowship funding through NC State University’s Genetics and Genomics Scholars, Molecular Biotechnology Training Grant (NIH T32GM133366) and the Genetics and Genomics Academy. We would also like to thank the NC State University’s Plant Transformation Facility for their help with tomato transformation, Genomic Sciences Laboratory for Sanger sequencing services, Cellular and Molecular Imaging Facility (NSF grant1624613) for providing user training and access to their equipment, Drs. Marcela Rojas-Pierce and Robert Franks for allowing us to use their microscopes, and members of the Alonso-Stepanova lab for their critical comments and suggestions.

## Author Contributions

A.N.S., J.M.A. and E.Z. conceived the project; A.N.S. and J.M.A. designed the study; J.P.F.M., M.F., A.E.Y., M.N., H.D., A.J.M., R.C., A.K., M.R., and A.N.S. designed and carried out experiments, and analyzed data; C.Z. verified and maintained GoldenBraid DNA constructs, and trained personnel; V.L., D.O., V.D. and E.Z. performed computational analysis; J.P.F.M., M.F., A.E.Y., and M.N. prepared figures; A.N.S., J.M.A., J.T.A.I., A.G., and E.Z. supervised research; J.P.F.M., M.F., A.E.Y., J.M.A., and A.N.S. wrote the manuscript. All authors read and approved the manuscript.

## Supplementary Data files

**Supplementary Figure S1.** Sequence composition of two previously described synthetic ethylene-inducible promoters, *5xEBS* and *10x2EBS-S10*.

**Supplementary Figure S2.** Sequence composition of the new synthetic ethylene-inducible *EBSn* promoter and its inactive (mutated) version, *mEBSn*.

**Supplementary Figure S3.** The *EBSn:3xYPet* reporter is induced by ACC in three-day-old dark-grown Arabidopsis seedlings.

**Supplementary Figure S4**. The roots of an *EBSn:3xYPet* reporter line in Arabidopsis are sensitive to as little as 10 nM ACC.

**Supplementary Figure S5**. The *EBSn* promoter activity is not fully abolished in three-day-old dark-grown heterozygous *ein4* Arabidopsis mutant seedlings.

**Supplementary Figure S6.** Ethylene-independent basal fluorescence is observed in root tips of three-day-old dark-grown *ein2-5 EBSn:3xYPet* line A seedlings but not in that of line C seedlings in Arabidopsis.

**Supplementary Figure S7.** *EBSn:GUS* shows similar patterns of expression and ethylene inducibility in multiple independent T-DNA lines in Arabidopsis.

**Supplementary Figure S8.** The *EBSn:GUS* reporter in Arabidopsis seedlings is turned on within 4 hours of exposure to ethylene.

**Supplementary Figure S9.** *EBSn*-driven fluorescent reporters show reproducible ethylene-inducible expression patterns in three-day-old dark-grown Arabidopsis seedlings.

**Supplementary Figure S10.** The loss of GUS enzymatic activity in some tissues of *EBSn:GUS* and *EBSn:GUS-1xYPet* lines may be the result of protein aggregation.

**Supplementary Figure S11.** The *EBSn* promoter is induced by ACC and wounding in Arabidopsis rosette leaves.

**Supplementary Figure S12.** Tomato leaf explants of independent *EBSn:GUS* and *EBSn:3xYPet* T0 lines exposed to ACC show varying levels of reporter activity.

**Supplementary Figure S13**. Tomato leaf explants of segregating T1 populations of *EBSn* reporter lines show expression level differences in ACC.

**Supplementary Figure S14.** Tomato backcrossing strategy enabled separation of multi-insertional events, elimination of transgene silencing, and identification of homozygous single-event *EBSn* lines.

**Supplementary Table 1.** GB clones harboring *EBSn* and *mEBSn* promoters for the ethylene-responsive fluorescent and histochemical reporters.

**Supplementary Table S2.** Segregation ratios and silencing events of tomato *EBSn* lines and their backcrossed offspring.

**Supplementary Table S3.** 14 ethylene-inducible, EIN3-targeted Arabidopsis thaliana genes that together harbor 18 *2EBS(−1)* elements in their promoters utilized in the construction of *EBSn*.

**Supplemental Table S4.** Nucleotide sequences for *EBSn* and *mEBSn* promoters and oligos

## References

1. Abeles FB, Morgan PW, Saltveit Jr ME. Ethylene (2nd ed.). In: Abeles FB, Morgan PW, Saltveit Jr ME, editors. Plant Biology, Academic Press, NY: Elsevier Inc. US; 1992. https://shop.elsevier.com/books/ethylene-in-plant-biology/abeles/978-0-08-091628-6

2. Alexander L, Grierson D. Ethylene biosynthesis and action in tomato: a model for climacteric fruit ripening. J Exp Bot. 2002:53(377):2039–2055. 10.1093/jxb/erf072

3. Almoguera C, Rojas A, Jordano J. Reversible Heat-induced inactivation of chimeric β-glucuronidase in transgenic plants. Plant Phys. 2002:129(1):333–341. 10.1104/pp.000992

4. Alonso JM, Hirayama T, Roman G, Nourizadeh S, Ecker JR. *EIN2*, a bifunctional transducer of ethylene and stress responses in Arabidopsis. Science. 1999:284(5423):2148-2152. 10.1126/science.284.5423.2148

5. Alonso JM, Stepanova AN, Solano R, Wisman E, Ferrari S, Ausubel FM, Ecker JR. Five components of the ethylene-response pathway identified in a screen for weak ethylene-insensitive mutants in Arabidopsis. Proc Natl Acad Sci U S A. 2003:100(5):2992–2997. 10.1073/pnas.0438070100

6. Alonso JM, Stepanova AN. A Recombineering-Based Gene Tagging System for Arabidopsis. In: Narayanan K, editors. Bacterial Artificial Chromosomes. Methods in Molecular Biology, vol 1227. Humana Press, New York, NY. 2015. pp 233-243. 10.1007/978-1-4939-1652-8_11

7. Althiab-Almasaud R, Sallanon H, Chang C, Chervin C. 1-Aminocyclopropane-1-carboxylic acid stimulates tomato pollen tube growth independently of ethylene receptors. Physiol Plant. 2021:173(4):1–7. 10.1111/ppl.13579

8. Binder BM. Ethylene signaling in plants. J Biol Chem. 2020:295(22):7710–7725. 10.1074/jbc.rev120.010854

9. Brumos J, Zhao C, Gong Y, Soriano D, Patel AP, Perez-Amador MA, Stepanova AN, Alonso JM. An improved recombineering toolset for plants. Plant Cell. 2020:32(1):100–122. 10.1105/tpc.19.00431

10. Chang C, Kwok SF, Bleecker AB, Meyerowitz EM. Arabidopsis ethylene-response gene ETR1: Similarity of Product to Two-Component Regulators. Science. 1993:262(5133):539-544. 10.1126/science.8211181

11. Chang KN, Zhong S, Weirauch MT, Hon G, Pelizzola M, Li H, Huang SSC, Schmitz RJ, Urich MA, Kuo D, et al. Temporal transcriptional response to ethylene gas drives growth hormone cross-regulation in Arabidopsis. Elife. 2013:11(2):e00675. 10.7554/elife.00675

12. Chen X, Zaro JL, Shen WC. Fusion protein linkers: Property, design and functionality. Adv Drug Deliv Rev. 2013:65(19):1357–1369. 10.1016/j.addr.2012.09.039

13. Clough SJ, Bent AF. Floral dip: a simplified method for Agrobacterium-mediated transformation of *Arabidopsis thaliana*. Plant J. 1998:16(6):735–743. 10.1046/j.1365-313x.1998.00343.x

14. Di Fino LM, Anjam MS, Besten M, Mentzelopoulou A, Papadakis V, Zahid N, Baez LA, Trozzi N, Majda M, Ma X, et al. Cellular damage triggers mechano-chemical control of cell wall dynamics and patterned cell divisions in plant healing. Dev Cell. 2025:60:1–12. 10.1016/j.devcel.2024.12.032

15. Dolgikh VA, Pukhovaya EM, Zemlyanskaya EV. Shaping ethylene response: the role of EIN3/EIL1 transcription factors. Front Plant Sci. 2019:10:1030. 10.3389/fpls.2019.01030

16. Eudes A, Mouille G, Thévenin J, Goyallon A, Minic Z, Jouanin L. Purification, cloning and functional characterization of an endogenous beta-glucoronidase in *Arabidopsis thaliana*. Plant Cell Physiol. 2008:49(9):1331–1341. 10.1093/pcp/pcn108

17. Fernandez-Moreno J-P, Stepanova AN. Monitoring ethylene in plants: Genetically encoded reporters and biosensors. Small Methods. 2019:4:1900260. 10.1002/smtd.201900260

18. Fernandez-Moreno J-P, Yaschenko AE, Neubauer M, Marchi AJ, Zhao C, Ascencio-Ibanez JT, Alonso JM, Stepanova AN. A rapid and scalable approach to build synthetic repetitive hormone-responsive promoters. Plant Biotech J. 2024:22:1942–1956. DOI: 10.1111/pbi.14313

19. Franck A, Guilley H, Jonard G, Richards K, Hirth L. Nucleotide sequence of Cauliflower Mosaic Virus DNA. Cell. 1980:21(1):285–294. 10.1016/0092-8674(80)90136-1

20. Gao Z, Wen C-K, Binder BM, Chen Y-F, Chang J, Chiang Y-H, Kerris III JK, Chang C, Schaller GE. Heteromeric interactions among ethylene receptors mediate signaling in Arabidopsis. J Biol Chem. 2008:283(35):23801–23810. 10.1074/jbc.m800641200

21. Hicks GR, Raikhel NV. Specific binding of nuclear localization sequences to plant nuclei. Plant Cell. 1993:5(8):983–994. 10.1105/tpc.5.8.983

22. Hua J, Chang C, Sun Q, Meyerowitz EM. Ethylene insensitivity conferred by Arabidopsis *ERS* gene. Science. 1995:269(5231):1712-1714. 10.1126/science.7569898

23. Hua J, Sakai H, Nourizadeh S, Chen QG, Bleecker AB, Ecker JR, Meyerowitz EM. *EIN4* and *ERS2* Are members of the putative ethylene receptor gene family in Arabidopsis. Plant Cell. 1998:10(8):1321–1332. 10.1105/tpc.10.8.1321

24. Jefferson RA, Kavanagh TA, Bevan MW. GUS fusions: beta-glucuronidase as a sensitive and versatile gene fusion marker in higher plants. EMBO J. 1987:6(13):3901–3907. 10.1002/j.1460-2075.1987.tb02730.x.

25. Johnson PR, Ecker JR. The ethylene gas signal transduction pathway: A molecular perspective. Annu Rev Genet. 1998:32:227–254. 10.1146/annurev.genet.32.1.227

26. Ju C, Yoon GM, Shemansky JM, Lin DY, Ying ZI, Chang J, Garrett WM, Kessenbrock M, Groth G, Tucker ML, et al. CTR1 phosphorylates the central regulator EIN2 to control ethylene hormone signaling from the ER membrane to the nucleus in Arabidopsis. Proc Natl Acad Sci U S A. 2012:109(47):19486–19491. 10.1073/pnas.1214848109

27. Jupe F, Rivkin AC, Michael TP, Zander M, Motley ST, Sandoval JP, Slotkin RK, Chen H, Castanon R, Nery JR, et al. The complex architecture and epigenomic impact of plant T-DNA insertions. PLoS Genet. 2019:15(1):e1007819. 10.1371/journal.pgen.1007819.

28. Kavita P, Burma PK. A comparative analysis of green fluorescent protein and beta-glucuronidase protein-encoding genes as a reporter system for studying the temporal expression profiles of promoters. J. Biosci. 2008:33(3):337–343. 10.1007/s12038-008-0053-4.

29. Kieber JJ, Rothenberg M, Roman G, Feldmann KA, Ecker JR. *CTR1*, a negative regulator of the ethylene response pathway in Arabidopsis, encodes a member of the raf family of protein kinases. Cell. 1993:72(3):427–441. 10.1016/0092-8674(93)90119-b

30. Kim J, Patterson SE, Binder BM. Reducing jasmonic acid levels causes *ein2* mutants to become ethylene responsive. FEBS Lett. 2013:587(2):226–230. 10.1016/j.febslet.2012.11.030

31. Klein JS, Jiang S, Galimidi RP, Keeffe JR, Bjorkman PJ. Design and characterization of structured protein linkers with differing flexibilities. Protein Eng Des Sel. 2014:27(10):325–330. 10.1093/protein/gzu043.

32. Krasnyanski SF, Sandhu J, Domier LL, Buetow DE, Korban SS. Effect of an enhanced CaMV 35S promoter and a fruit-specific promoter on *uida* gene expression in transgenic tomato plants. In Vitro Cell Dev Biol Plant. 2001:37:427–433. 10.1007/s11627-001-0075-1

33. Kurihara D, Mizuta Y, Sato Y, Higashiyama T. ClearSee: a rapid optical clearing reagent for whole-plant fluorescence imaging. Development. 2015:142(23):4168–79. 10.1242/dev.127613

34. Li W, Ma M, Feng Y, Li H, Wang Y, Ma Y, Li M, An F, Guo H. *EIN2*-directed translational regulation of ethylene signaling in Arabidopsis. Cell. 2015:163(3):670–683. 10.1016/j.cell.2015.09.037

35. Li S, Zhu B, Pirrello J, Xu C, Zhang B, Bouzayen M, Chen K, Grierson D. Roles of RIN and ethylene in tomato fruit ripening and ripening-associated traits. New Phytol. 2019:226(2):460–475. 10.1111/nph.16362

36. Li D, Flores-Sandoval E, Ahtesham U, Coleman A, Clay JM, Bowman JL, Chang C. Ethylene-independent functions of the ethylene precursor ACC in Marchantia polymorpha. Nature Plants. 2020:6(11):1335–1344. 10.1038/s41477-020-00784-y

37. Li D, Mou W, Van de Poel B, Chang C. Something old, something new: Conservation of the ethylene precursor 1-amino-cyclopropane-1-carboxylic acid as a signaling molecule. Curr Opin Plant Biol. 2022:65:102116. 10.1016/j.pbi.2021.102116

38. Mao K, Zhang M, Kong Y, Dai S, Wang Y, Meng Q, Ma N, Lv W. Origin, expansion, and divergence of ETHYLENE-INSENSITIVE 3 (EIN3)/EIN3-LIKE transcription factors during streptophytes evolution. Front Plant Sci. 2022:13:858477. 10.3389/fpls.2022.858477

39. Mattoo AK (1991) The Plant Hormone Ethylene (1st ed.). In Mattoo AK and Suttle JC editors. CRC Press, Boca Raton, FL: US. 1991. p. 347. 10.1201/9781351075763

40. Matsuura T, Hosoda K, Ichihashi N, Kazuta Y, Yomo T. Kinetic Analysis of *β*-Galactosidase and *β*-Glucuronidase tetramerization coupled with protein translation. J Biol Chem. 2011:286(25):22028–22034. 10.1074/jbc.m111.240168

41. Merchante C, Brumos J, Yun J, Hu Q, Spencer KR, Enríquez P, Binder BM, Heber S, Stepanova AN, Alonso JM. gene-specific translation regulation mediated by the hormone-signaling molecule *EIN2*. Cell. 2015:163(3):684–697. 10.1016/j.cell.2015.09.036

42. Mlotshwa S, Pruss GJ, Gao Z, Mgutshini NL, Li J, Chen X, Bowman LH, Vance V. Transcriptional silencing induced by Arabidopsis T-DNA mutants is associated with 35S promoter siRNAs and requires genes involved in siRNA-mediated chromatin silencing. Plant J. 2011:64:699–704. 10.1111/j.1365-313X.2010.04358.x.

43. Mou W, Kao Y-T, Michard E, Simon AA, Li D, Wudick MM, Lizzio MA, Feijó JA, Chang C. Ethylene-independent signaling by the ethylene precursor ACC in Arabidopsis ovular pollen tube attraction. Nature Commun. 2020:11:4082. 10.1038/s41467-020-17819-9

44. Mou W, Khare R, Polko JK, Taylor I, Xu J, Xue D, Benfey P, Van de Poel B, Chang C, Kieber JJ. Ethylene-independent modulation of root development by ACC via downregulation of WOX5 and group I CLE peptide expression. Proc Natl Acad Sci U S A. 2025:122(6):e2417735122. 10.1073/pnas.2417735122

45. Nguyen AW, Daugherty PS. Evolutionary optimization of fluorescent proteins for intracellular FRET. Nat Biotechnol. 2005:23:355–360. 10.1038/nbt1066

46. O’Malley RC, Huang SC, Song L, Lewsey MG, Bartlett A, Nery JR, Galli M, Gallavotti A, Ecker JR. Cistrome and epicistrome features shape the regulatory DNA landscape. Cell. 2016:165(5):1280–1292. 10.1016/j.cell.2016.04.038

47. Odell JT, Nagy F, Chua NH. Identification of DNA sequences required for activity of the cauliflower mosaic virus 35S promoter. Nature. 1985:313:810–812. 10.1038/313810a0

48. Pandey BK, Huang G, Bhosale R, Hartman S, Sturrock CJ, Jose L, Martin OC, Karady M, Voesenek LACJ, Ljung K, et al. Plant roots sense soil compaction through restricted ethylene diffusion. Science. 2021:371(6526):276-280. 10.1126/science.abf3013

49. Park HL, Seo DH, Lee HY, Bakshi A, Park C, Chien Y-C, Kieber JJ, Binder BM, Yoon GM. Ethylene-triggered subcellular trafficking of *CTR1* enhances the response to ethylene gas. Nat Commun. 2023:14:365. 10.1038/s41467-023-35975-6

50. Quinet M, Angosto T, Yuste-Lisbona FJ, Blanchard-Gros R, Bigot S, Martinez J-P, Lutts S. Tomato fruit development and metabolism. Front Plant Sci. 2019:10:1554. 10.3389/fpls.2019.01554

51. Sakai H, Hua J, Chen QG, Meyerowitz EM. *ETR2* is an *ETR1-like* gene involved in ethylene signaling in Arabidopsis. Proc Natl Acad Sci U S A. 1998:95(10):5812–5817. 10.1073/pnas.95.10.5812

52. Sarrion-Perdigones A, Vazquez-Vilar M, Palací J, Castelijns B, Forment J, Ziarsolo P, Blanca J, Granell A, Orzaez D. GoldenBraid 2.0: A comprehensive DNA assembly framework for plant synthetic biology. Plant Physiol. 2013:162(3):1618–1631. 10.1104/pp.113.217661

53. Schubert D, Lechtenberg B, Forsbach A, Gils M, Bahadur S, Schmidt R. Silencing in Arabidopsis T-DNA Transformants: The predominant role of a gene-specific RNA sensing mechanism versus position effects. Plant Cell. 2004:16(10):2561–2572. 10.1105/tpc.104.024547

54. Solano R, Stepanova AN, Chao Q, Ecker JR. Nuclear events in ethylene signaling: a transcriptional cascade mediated by ETHYLENE-INSENSITIVE3 and ETHYLENE-RESPONSE-FACTOR1. Genes Dev, 1998:12(23):3703–14. 10.1101/gad.12.23.3703

55. Song J, Zhu C, Zhang X, Wen X, Liu L, Peng J, Guo H, Yi C. Biochemical and structural insights into the mechanism of DNA recognition by Arabidopsis *ETHYLENE INSENSITIVE3*. PLoS One. 2015:10(9): e0137439. 10.1371/journal.pone.0137439

56. Stepanova AN. Nuclear events in ethylene signaling in *Arabidopsis thaliana*. PhD Thesis, University of Pennsylvania. 2001. Dissertation available from ProQuest AAI3015380.

57. Stepanova AN, Hoyt JM, Hamilton AA, Alonso JM. A Link between ethylene and auxin uncovered by the characterization of two root-specific ethylene-insensitive mutants in Arabidopsis. Plant Cell. 2005:17(8):2230–42. 10.1105/tpc.105.033365

58. Stepanova AN, Yu J, Likhacheva AV, Alonso JM. Multilevel interactions between ethylene and auxin in Arabidopsis roots. Plant Cell. 2007:19:2169–2185. 10.1105/tpc.107.052068

59. Tsukanov AV, Mironova VV, Levitsky VG. Motif models proposing independent and interdependent impacts of nucleotides are related to high and low affinity transcription factor binding sites in Arabidopsis. Front Plant Sci. 2022:13:938545. 10.3389/fpls.2022.938545

60. Eck JV, Keen P, Tjahjadi M. Agrobacterium tumefaciens-Mediated Transformation of Tomato. In: Kumar S, Barone P, Smith M, editors. Transgenic Plants. Methods in Molecular Biology, vol 1864. Humana Press, New York, NY: USA. 2019. 10.1007/978-1-4939-8778-8_16

61. Vanderstraeten L, Depaepe T, Bertrand S, Van Der Straeten D. The ethylene precursor ACC affects early vegetative development independently of ethylene signaling. Front Plant Sci. 2019:10:1591. 10.3389/fpls.2019.01591

62. Wang R, Lammers M, Tikunov Y, Bovy AG, Angenent GC, de Maagd RA. The rin, *nor* and *Cnr* spontaneous mutations inhibit tomato fruit ripening in additive and epistatic manners. Plant Sci. 2020:294:110436. 10.1016/j.plantsci.2020.110436

63. Wang L, Ko EE, Tran J, Qiao H. TREE1-EIN3–mediated transcriptional repression inhibits shoot growth in response to ethylene. Proc Natl Acad Sci U S A. 2020:117(46):29178–29189. 10.1073/pnas.2018735117

64. Wang L, Zhang Z, Zhang F, Shao Z, Zhao B, Huang A, Tran J, Hernandez FV, Qiao H. EIN2-directed histone acetylation requires EIN3-mediated positive feedback regulation in response to ethylene. Plant Cell. 2021:33(2):322–337. 10.1093/plcell/koaa029

65. Wasserman WW, Sandelin A. Applied bioinformatics for the identification of regulatory elements. Nat Rev Genet. 2004:5:276–287. 10.1038/nrg1315

66. Wen X, Zhang C, Ji Y, Zhao Q, He W, An F, Jiang L, Guo H. Activation of ethylene signaling is mediated by nuclear translocation of the cleaved EIN2 carboxyl terminus. Cell Res. 2012:22:1613–1616. 10.1038/cr.2012.145

67. Zhao C, Yaschenko A, Alonso JM, Stepanova AN. Leveraging synthetic biology approaches in plant hormone research. Curr Opin Plant Biol. 2021:60:101998. 10.1016/j.pbi.2020.101998

68. Zhou R, Benavente LM, Stepanova AN, Alonso JM. A recombineering-based gene tagging system for Arabidopsis. Plant J. 2011:66:712–723. 10.1111/j.1365-313x.2011.04524.x

