## Supplementary data for "*EBSn,* a robust synthetic reporter for monitoring ethylene responses in plants"

### Supplementary Figures

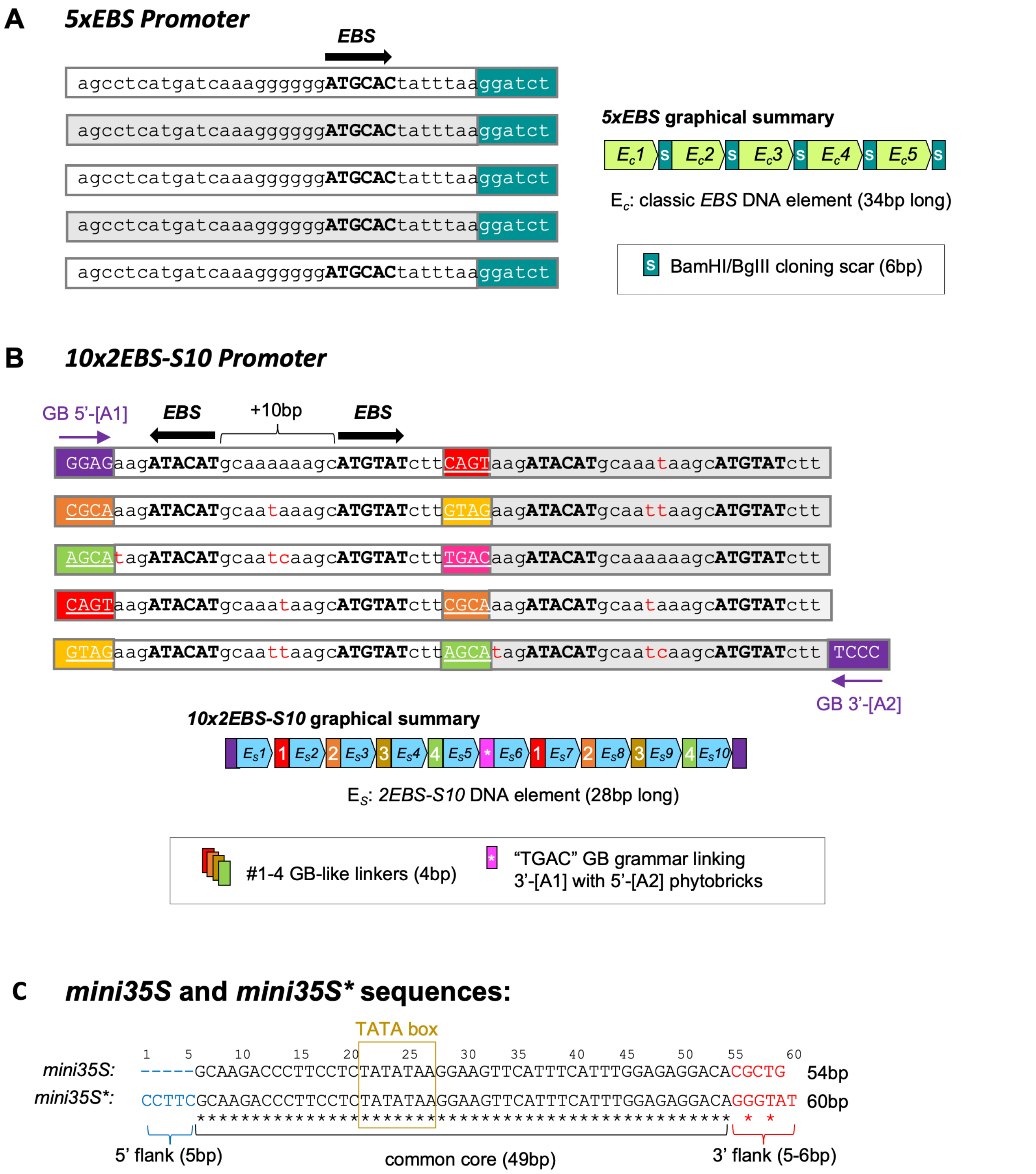


**Supplementary Figure S1. Sequence composition of two previously described synthetic ethylene-inducible promoters, *5xEBS* and *10x2EBS-S10*. A)** The *5xEBS* promoter

(Stepanova 2001; Stepanova at al., 2007) consists of five 34bp-long repeats (white and grey) placed in tandem, harboring a single forward-oriented *EBS* element each, and separated by 6-bp-long BamHI/BglII cloning scars (turquoise). **B)** The *10x2EBS-S10* promoter (Fernandez-Moreno et al., 2024) consists of ten 28-bp-long repeats (white and grey) placed in tandem and separated by 4-bp-long GB (TGAC, pink) or GB-like (1-CAGT red, 2-CGCA orange, 3-GTAG mustard, and 4-AGCA light green) grammar codes. Each repeat harbors a dual *2EBS-S10* element (Song et al. 2015) which consists of two everted sequences separated by a 10 nucleotide spacer (+10bp), with divergent nucleotides marked in red. The *10x2EBS-S10* promoter is flanked by 5'-[A1] and 3'-[A2] GB grammar codes, thus forming the final [A1-A2] distal promoter phytobrick (Fernandez-Moreno et al. 2024). **C)** Nucleotide sequence comparison of *mini35S* and *mini35S** core promoters. The alignment of *mini35S** (originally from the *5xEBS:GUS* reporter in Stepanova 2001; Stepanova at al., 2007; and from *10x2EBS-S10:GUS* in Fernandez-Moreno et al. 2024; herein used in the *EBSn:GUS* and *mEBSn:GUS* constructs to enable their comparisons with *5xEBS:GUS*) and *mini35S* (originally from the *10x2EBS-S10:3xYPet* reporter in Fernandez-Moreno et al. 2024; herein used in the *EBSn:3xYPet* and *EBSn:GUS-1xYPet* constructs, see **Fig. 1B**) sequences reveals a common TATA-box-containing 49bp core but divergent 5' and 3' flanks. *mini35S* lacks the 5-bp 5' sequence segment present in *mini35S** and has a different 5bp 3' flank relative to the 6bp 3' flank of *mini35S**.

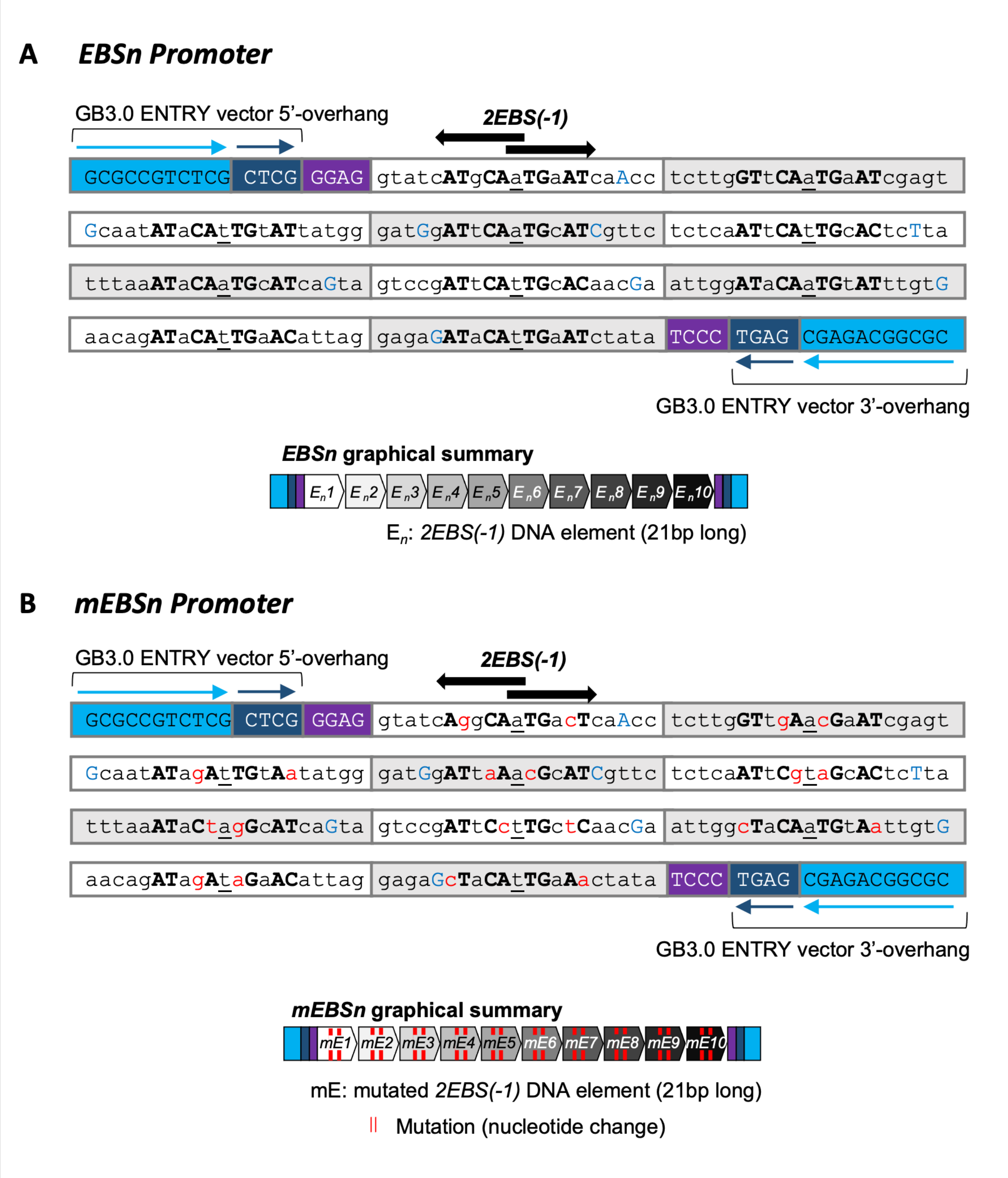


**Supplementary Figure S2. Sequence composition of the new synthetic ethylene-inducible *EBSn* promoter and its inactive (mutated) version, *mEBSn*. A)** The *EBSn* promoter consists of ten 21-bp-long naturally occurring, non-identical repeats randomly stacked back to back without additional spacers and harboring two everted *EBS* elements (black arrows) each that overlap by a single nucleotide (underlined). For each *2EBS(-1)* element, nucleotides presumed to be essential for EIN3 binding are marked in bold black capital letters, whereas single-nucleotide substitutions introduced in the flanks of most 21bp elements to remove undesired possible binding sites of non-target transcription factors (distinct from the target EIN3) are shown in blue capital letters. **B)** The *mEBSn* promoter. To create the mutated version of *EBSn*, two additional nucleotide substitutions (red lowercase) were made, one per each *EBS* half-site, to disrupt their EIN3-binding ability. The *EBSn* and *mEBSn* promoters, similarly to the *10x2EBS-S10* promoter (**Supplementary Fig. S1B**) are flanked by 5'-[A1] and 3'-[A2] GB grammar codes, producing the final [A1-A2] distal promoter phytobricks (**Supplementary Table S1A**). Overhangs for cloning both promoters into GB entry-level (*pUPD2* vector) were added to both flanks during the domestication. They consist of a type IIS BsmBI restriction site (cyan) and a *pUPD2* vector-compatible 4bp grammar code (navy blue).

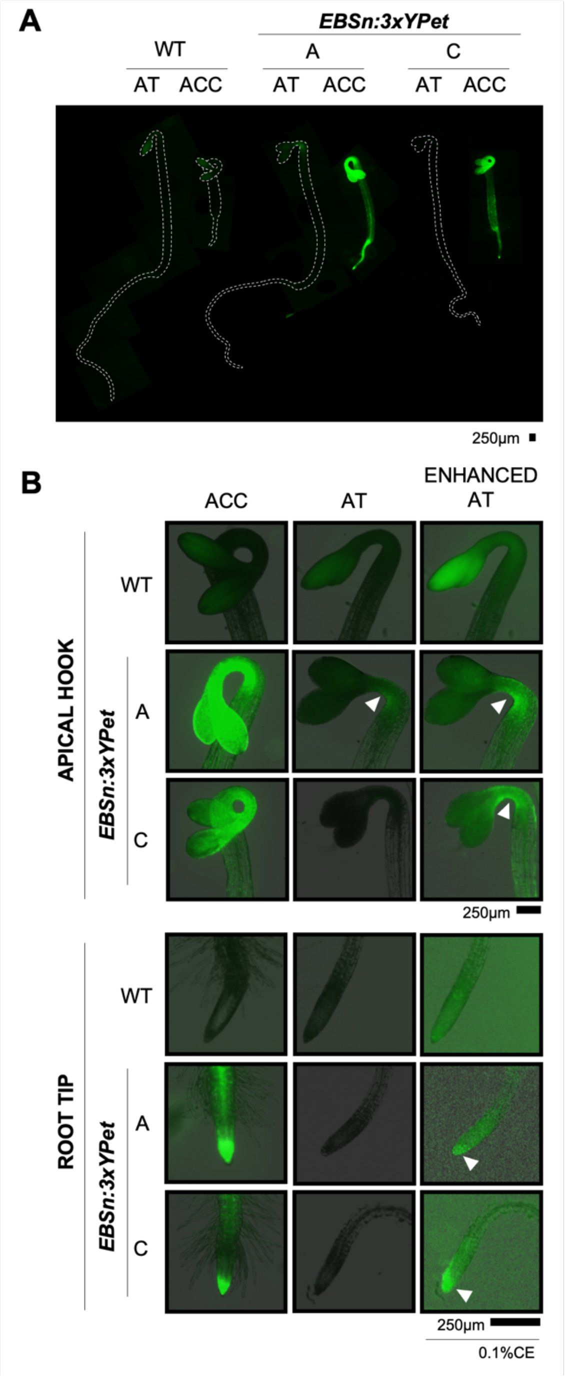


**Supplementary Figure S3. The *EBSn:3xYPet* reporter is induced by ACC in three-day-old dark-grown Arabidopsis seedlings. A)** Whole-plant images of wild-type (WT) and *EBSn:3xYPet* line A and C seedlings germinated in plain AT media or AT media supplemented with 10 µm ACC (ACC). **B)** Close-up apical hook and root tip images of seedlings from panel A. Arrowheads point to weak hook fluorescence observed in the nuclei of *EBSn:3xYPet* line A grown in AT media but not in line C or wild-type Col-0 plants (WT). Digital enhancement of AT images (0.1% contrast enhancement, CE, in ImageJ v1.52, Fiji 2015) reveals nuclear localization of fluorescence in the apical hooks of both *EBSn:3xYPet* lines (arrowheads), as well as some fluorescence signal in transgenic root tips (arrowheads), but not in WT.

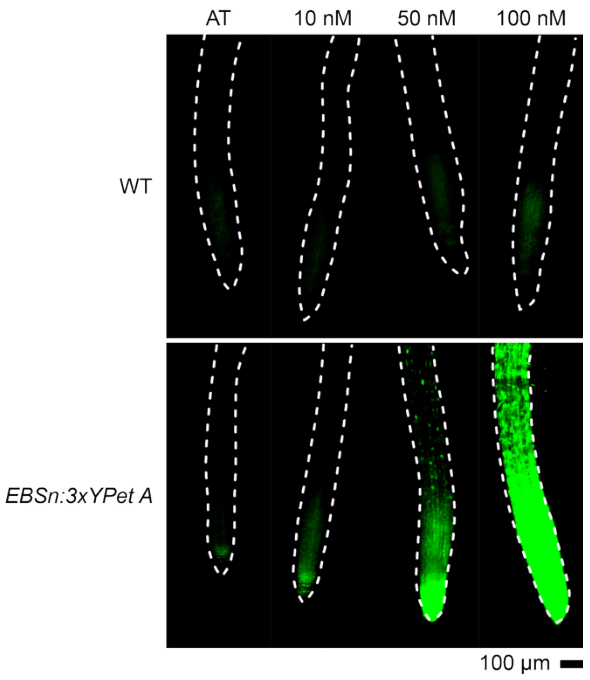


**Supplementary Figure S4**. **The roots of an *EBSn:3xYPet* reporter line in Arabidopsis are sensitive to as little as 10 nM ACC.** Fluorescence of primary roots of 10-day-old line A seedlings grown vertically under continuous light in plates supplemented with the indicated concentrations of ACC are shown.

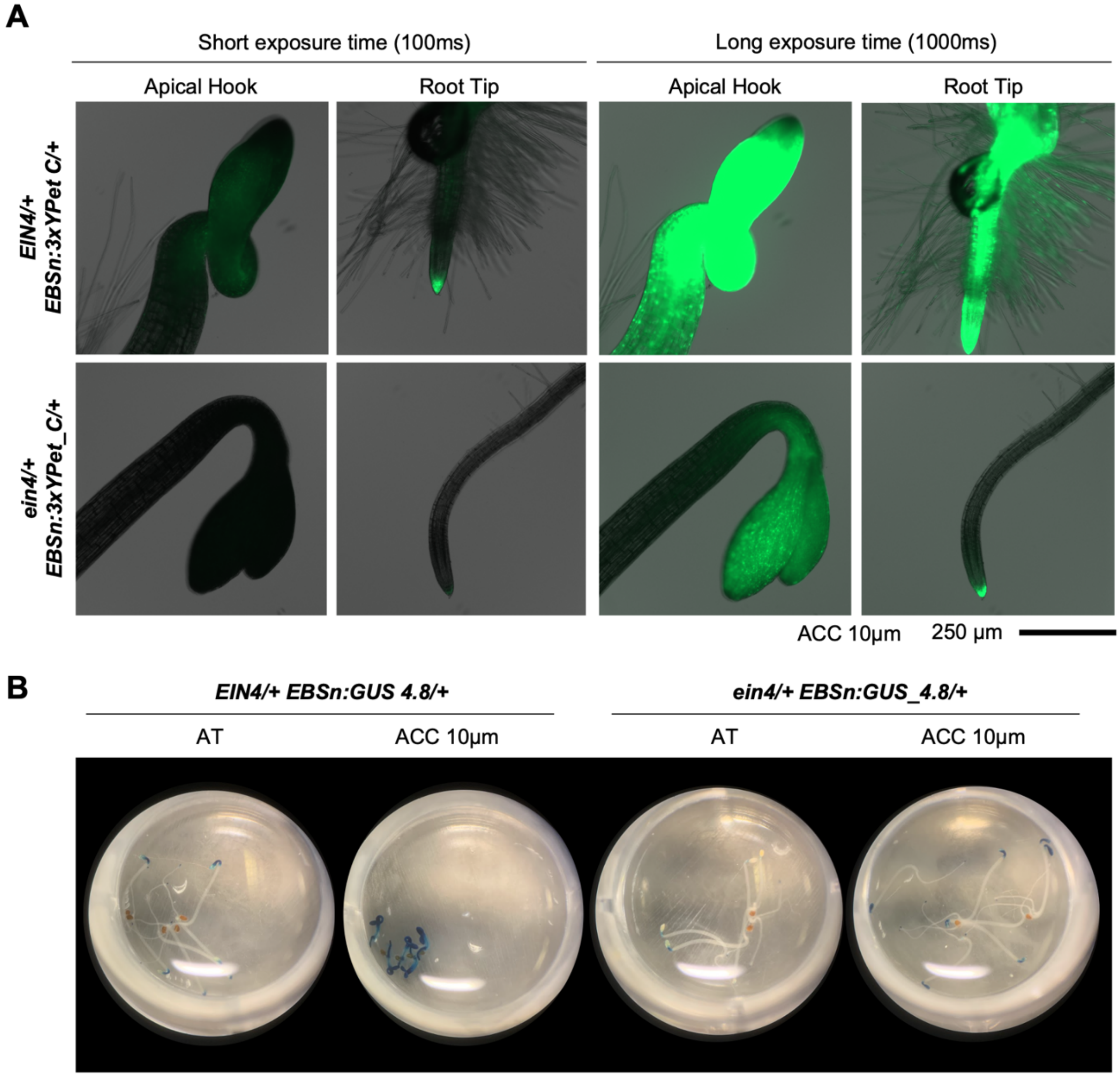


**Supplementary Figure S5**. **The *EBSn* promoter activity is not fully abolished in three-day-old, dark-grown, heterozygous *ein4* Arabidopsis mutant seedlings. A)** 3xYpet fluorescence of *EBSn:3xYPet/*+ and *ein4/*+ *EBSn:3xYPet/*+ F1 seedlings germinated in media supplemented with 10 µm ACC (line C). **B)** GUS staining of *EBSn:GUS/*+ and *ein4/*+ *EBSn:GUS/*+ F1 seedlings germinated in plain AT media or AT media supplemented with 10 µm ACC (line 4.8). Note the residual ethylene inducibility of the *EBSn:GUS* and *EBSn:3xYPet* reporters in *ein4* heterozygotes.

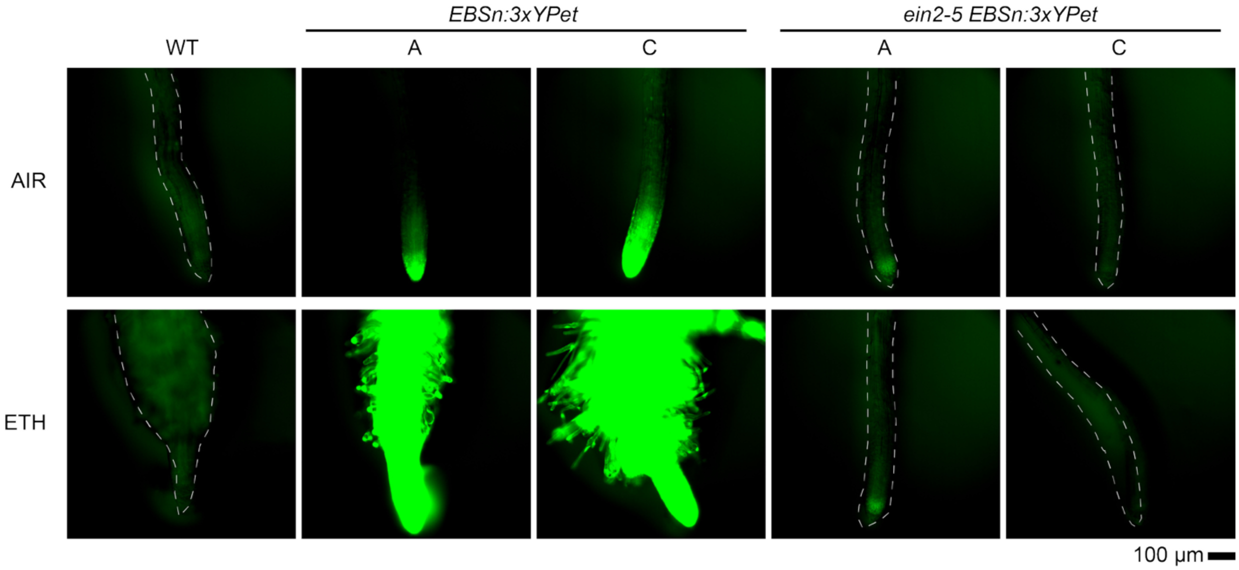


**Supplementary Figure S6. Ethylene-independent basal fluorescence is observed in root tips of three-day-old dark-grown *ein2-5 EBSn:3xYPet* line A seedlings but not in that of line C seedlings in Arabidopsis.** Seeds were germinated in AT plates in the presence of hydrocarbon-free air (AIR) or 10 ppm ethylene (ETH). Pictures were digitally enhanced for the purpose of showing *ein2-5 EBSn:3xYPet* line A activity. ETH = 10 ppm ethylene.

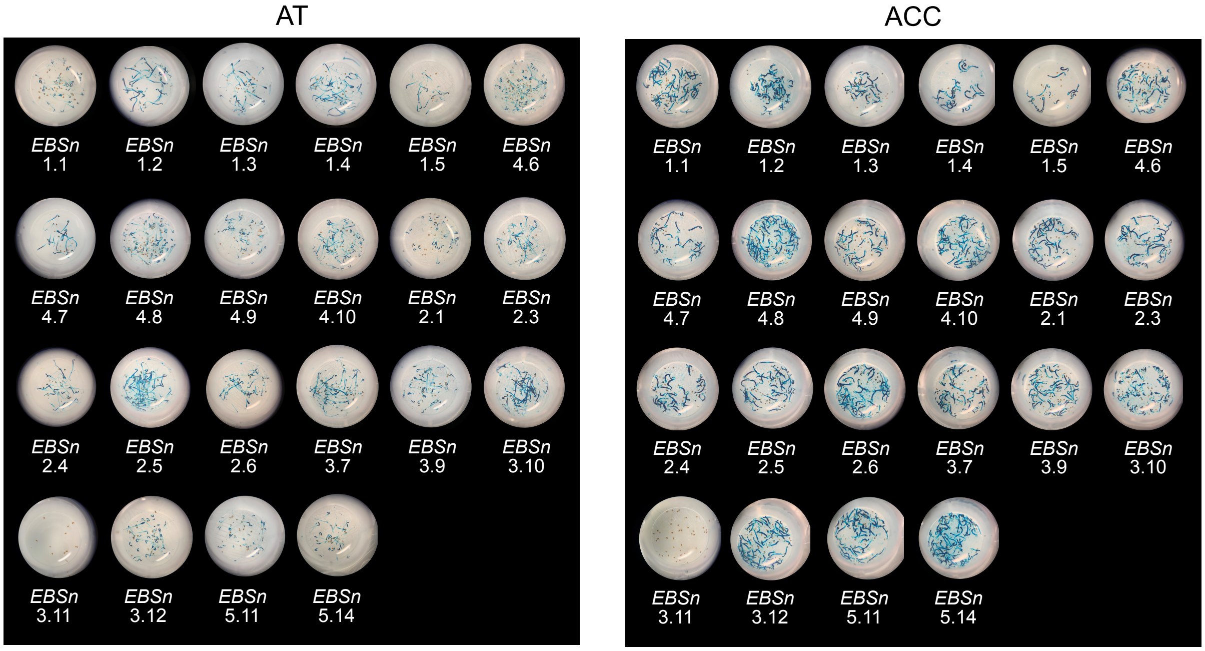


**Supplementary Figure S7. *EBSn:GUS* shows similar patterns of expression and ethylene inducibility in multiple independent T-DNA lines in Arabidopsis.** Three-day-old etiolated *EBSn:GUS* T2 seedlings grown either in AT media or AT media supplemented with 10 μM ACC were stained for GUS overnight.  Line 3.11 turned out to be sensitive to Basta and serves as a negative control.

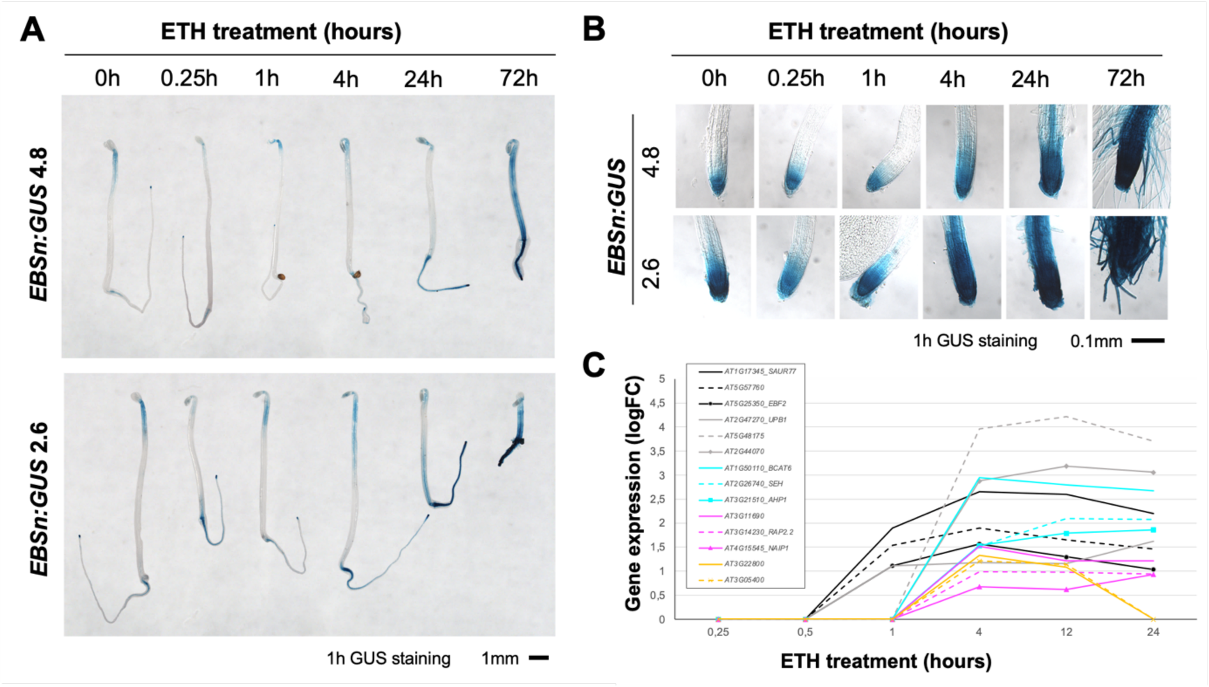


**Supplementary Figure S8. The *EBSn:GUS* reporter in Arabidopsis seedlings is turned on within 4 hours of exposure to ethylene.** The induction timing of *EBSn* is consistent with the ethylene induction kinetics of genes whose *cis*-elements were utilized in the *EBSn* promoter design. **A)** Three-day-old dark-grown *EBSn:GUS* seedlings (lines 4.8 and 2.6) germinated in AT plates incubated in hydrocarbon-free air (AIR) or 10 ppm ethylene (ETH). Plates with seedlings were moved to ethylene for the indicated amount of time, and samples were harvested when seedlings were 72-hour-old. GUS staining was performed only for 1 hour to avoid reaction oversaturation. **B)** Root tips of representative three-day-old dark-grown GUS-stained *EBSn:GUS* seedlings (lines 4.8 and 2.6) from the same populations as shown in panel A. **C)** Ethylene induction kinetics of 14 genes whose promoters served as a source of *2EBS(-1)* elements utilized in the design of *EBSn*. Log2FC values were retrieved from an RNA-seq time-course experiment reported by Chang et al. 2013. Only statistically significant log2FC values (at FDR < 0.05) are depicted.

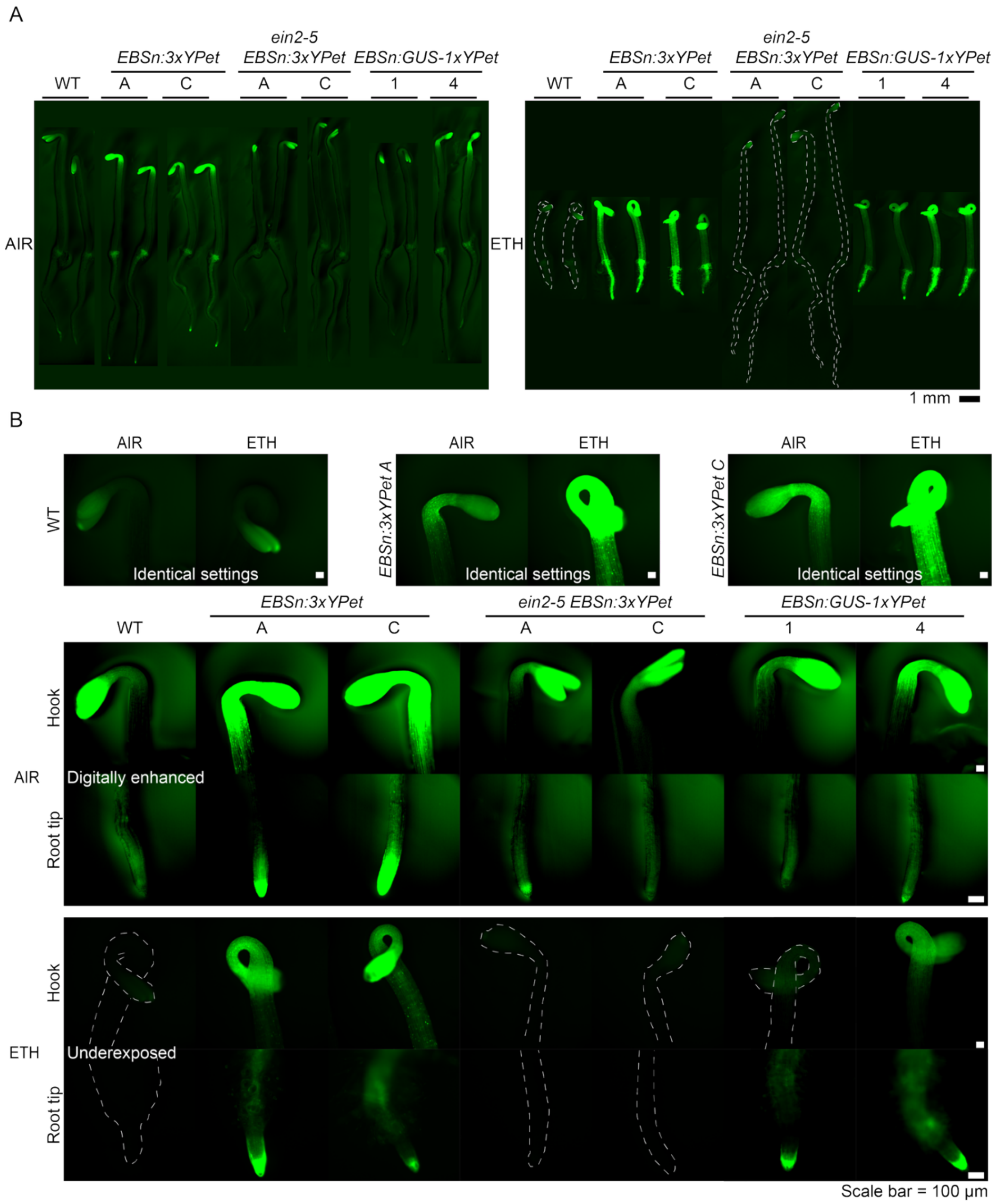


**Supplementary Figure S9. *EBSn*-driven fluorescent reporters show reproducible ethylene-inducible expression patterns in three-day-old dark-grown Arabidopsis seedlings. A)** Whole-plant images of wild-type (WT) and transgenic seedlings grown in AT plates in the presence of hydrocarbon-free air (AIR) or 10 ppm ethylene (ETH). **B)** Close-up images of representative apical hooks and roots from the populations shown in panel A. The root images displayed in panel B are the same as shown in Supplementary Figure S6 with a different contrast/brightness setting. Note that all AIR (but not ETH) pictures in panel B were digitally enhanced to show detectable activity in *EBSn:GUS-1xYPet* lines in air. Thus, the expression levels in panel B are comparable within a treatment (AIR or ETH), but not among treatments, with the ETH images intentionally underexposed to avoid fluorescence signal oversaturation.

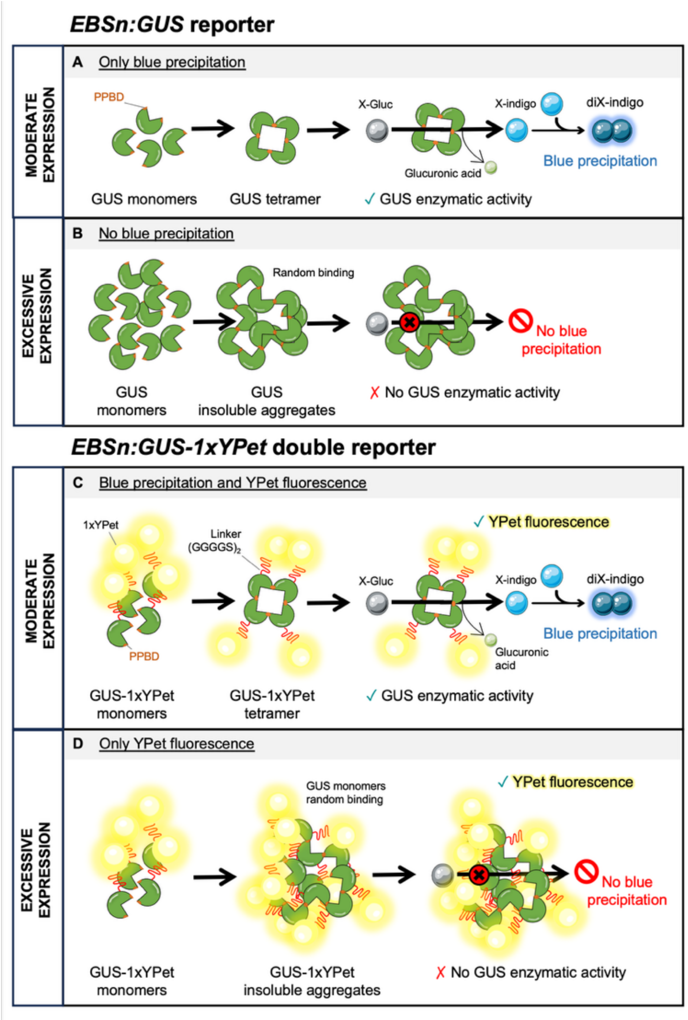


**Supplementary Figure S10. The loss of GUS enzymatic activity in some tissues of *EBSn:GUS* and *EBSn:GUS-1xYPet* lines may be the result of protein aggregation. A)** Low to moderate levels of *EBSn:GUS* expression result in the production of functional GUS (beta-glucuronidase) enzyme tetramers that can metabolize the X-Gluc precursor into glucuronic acid and X-indigo, leading to blue tissue staining upon the dimerization of two X-indigo molecules. GUS monomers and dimers are known to be enzymatically inactive, but become activated upon assembly into tetramers (Matsuura et al. 2011). **B)** Excessive levels of *EBSn:GUS* expression may lead to the loss of GUS activity and, therefore, of GUS staining. We speculate that overproduction of GUS enzymes may trigger GUS protein aggregation, impeding the access of GUS to the X-Gluc substrate and abolishing the production of diX-indigo blue precipitate. Since GUS tetramers form via protein-protein interactions, we speculate that when expressed at excessive levels, GUS subunits assemble randomly, generating insoluble, non-functional GUS aggregates. **C)** Low to moderate expression levels of the *EBSn:GUS-1xYPet* dual reporter result in functional GUS activity and 1xYPet fluorescence of the GUS-YPet fusion protein. **D)** Excessive levels of the *EBSn:GUS-1xYPet* dual reporter expression result in the loss of GUS activity and no blue staining despite maintaining functional 1xYPet. We hypothesize that GUS aggregation would not interfere with proper 1xYPet folding, thus preserving 1xYPet fluorescence in GUS-1xYPet aggregates. PPBD = protein-protein binding domain. Image icons were obtained from Bioicons (https://bioicons.com).

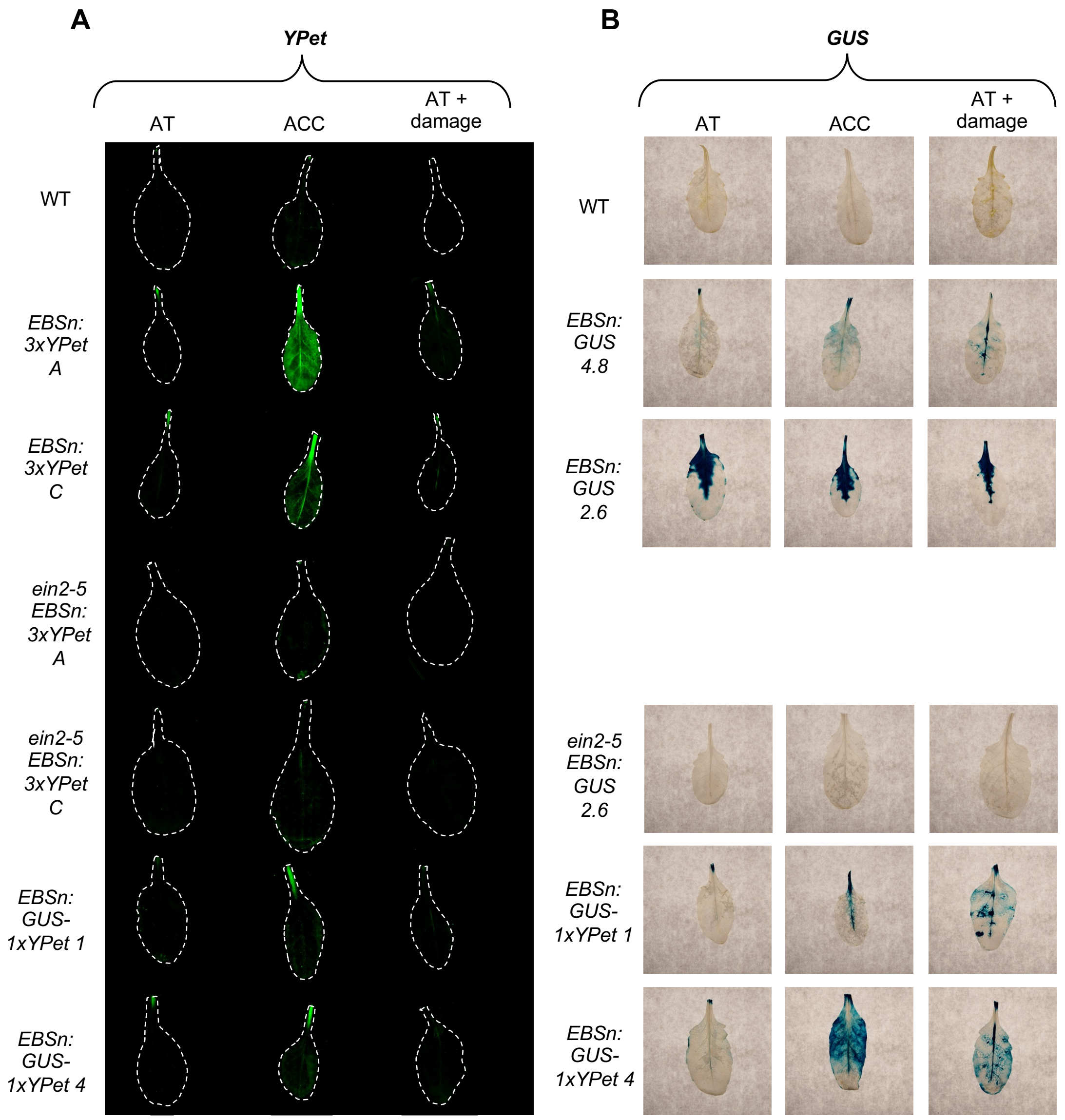


**Supplementary Figure S11. The *EBSn* promoter is induced by ACC and wounding in Arabidopsis rosette leaves.** Detached leaves of 45-day-old plants were incubated for 24 hours in plain AT and AT supplemented with 10 μM ACC, or damaged by gentle forcep smashing followed by a 24-hour incubation in plain AT. **A)** YPet fluorescence. **B)** GUS staining.

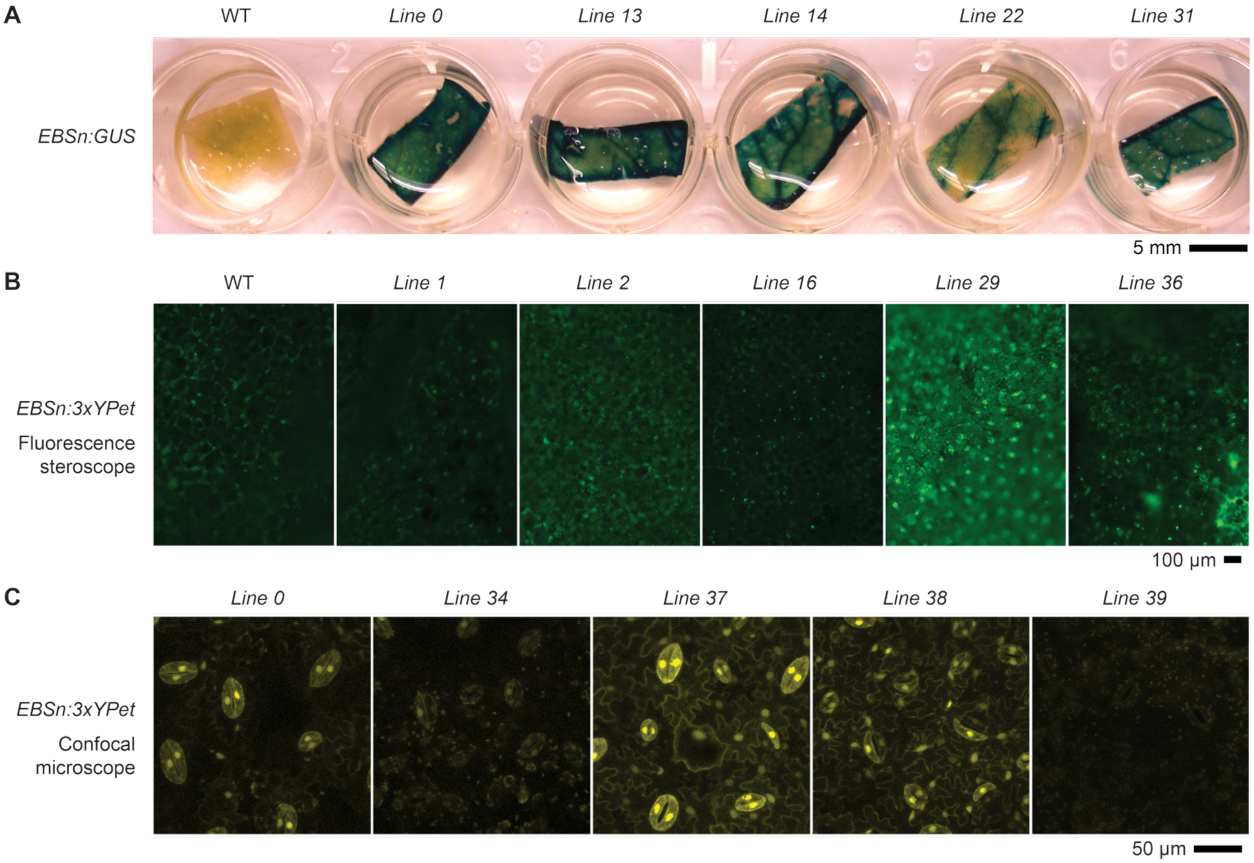


**Supplementary Figure S12. Tomato leaf explants of independent *EBSn:GUS* and *EBSn:3xYPet* T0 lines exposed to ACC show varying levels of reporter activity**. All explants were treated with 50 µM ACC for 24 hours. **A)** GUS staining of *EBSn:GUS* leaf sectors. Five independent T0 lines are shown. **B), C)** 3xYpet fluorescence in *EBSn:3xYPet* leaf sectors. The first five T0 lines were screened by fluorescence stereoscope (panel B), and the next five T0 lines were examined using ,  microscopy to better visualize the 3xYPet fluorescence in stomatal guard cells (panel C).

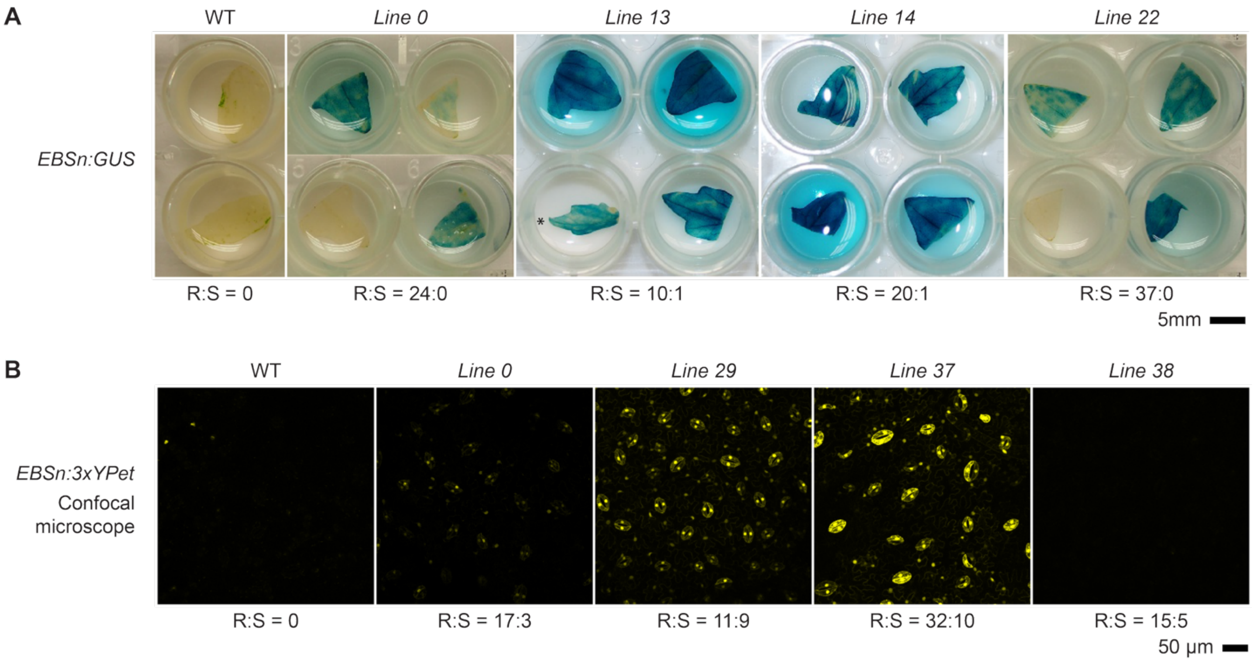


**Supplementary Figure S13**. **Tomato leaf explants of segregating T1 populations of *EBSn* reporter lines show expression level differences in ACC.** Transgenic plants were germinated and grown in AT medium supplemented with 100 mg/L kanamycin. WT plants (*Solanum lycopersicum* M82) were germinated and grown in plain AT. Leaf explants were treated with 50 µM ACC for 24 hours. **A)** GUS staining of *EBSn:GUS* leaf sectors. Note that some silencing is evident, with some leaves from kanamycin-resistant plants not staining for GUS (note white explants in Line 0 and Line 22) and leaves from kanamycin-sensitive plants being positive for GUS (note blue explants marked with * in Line 13). **B**) 3xYPet fluorescence of *EBSn:3xYPet* leaf sectors. Note that the reporter activity declined in Line 0 and Line 38 in T1. Among the segregating populations, *EBSn:GUS* lines 13 and 14 and *EBSn:3xYPet* lines 29 and Line 37 showed homogenous patterns of *EBSn* reporter expression across multiple siblings and similar levels between lines. Kanamycin resistant-to-sensitive (R:S) plant ratios suggested more than one insertional event in all of the lines tested, necessitating backcrosses (see Supplementary Table 2).

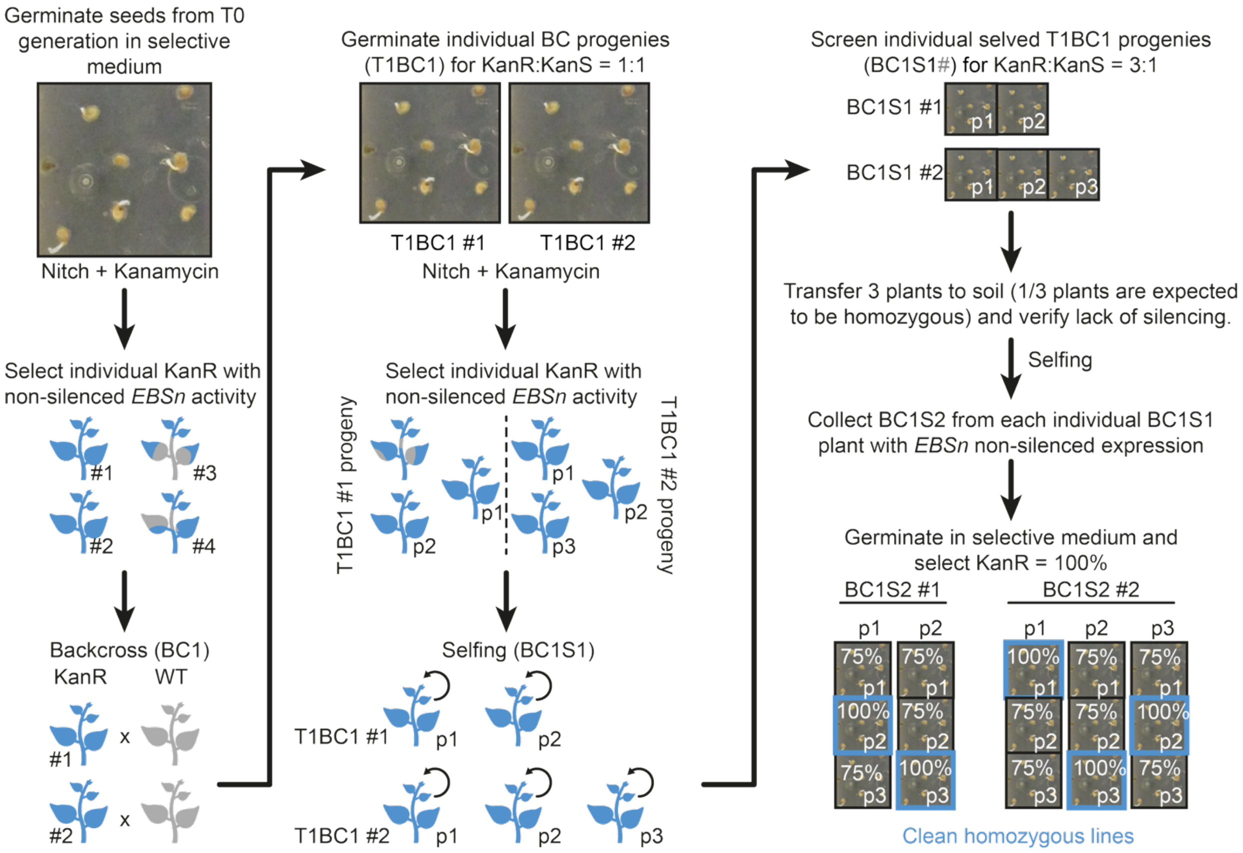


**Supplementary Figure S14. Tomato backcrossing strategy enabled separation of multi-insertional events, elimination of transgene silencing, and identification of homozygous single-event *EBSn* lines**. KanR and KanS are kanamycin-resistant and -sensitive plants, respectively. #1 and #2 refer to two siblings that are independently backcrossed to WT (M82). p1-p3 are the independent progeny of the T1BC1 population.

### Supplementary Data Sets

**Supplementary Table 1**

https://docs.google.com/spreadsheets/d/1R1zA3s59BPwXhBbnqJqA6hEdsH7yFAHr/edit?gid=1032736132#gid=1032736132

**Supplementary Table 2**

https://docs.google.com/spreadsheets/d/1iyM86s3bjUcYRPyrsVC0ellLx0kbYFyb/edit?gid=2104734906#gid=2104734906

**Supplementary Table 3**

https://docs.google.com/spreadsheets/d/1kv04fIRCA2FG0WMLRJRoFpBqLqZsU5_8/edit?gid=1819316199#gid=1819316199

**Supplementary Table 4**

https://docs.google.com/spreadsheets/d/1eYchSXzdqzyqbHKHTrTrICbrq_25YanP/edit?gid=2003281990#gid=2003281990
